# *O*-GlcNAcylation of Keratin 18 coordinates TCA cycle to promote cholangiocarcinoma progression

**DOI:** 10.1101/2023.07.23.550247

**Authors:** Xiangfeng Meng, Yue Zhou, Lei Xu, Changjiang Wang, Xiao Tian, Xiang Zhang, Yi Hao, Bo Cheng, Lei Wang, Jialin Liu, Ran Xie

**Affiliations:** State Key Laboratory of Coordination Chemistry, School of Chemistry and Chemical Engineering, Chemistry and Biomedicine Innovation Center (ChemBIC), Nanjing University, Nanjing, China; Department of Gastroenterology, Nanjing Drum Tower Hospital, The Affiliated, Hospital of Nanjing University Medical School, Nanjing, China; State Key Laboratory of Proteomics, Beijing Proteome Research Center, National Center for Protein Sciences (Beijing), Beijing Institute of Lifeomics, Beijing, China; College of Chemistry and Molecular Engineering, Peking University, Beijing, China; School of Pharmaceutical Sciences, Peking University, Beijing, China

## Abstract

Glycosylation in human cholangiocarcinoma (CCA) actively contributes to pathophysiological steps of tumor progression. Of note is the dynamic modification of proteins by *O*-linked β-*N*-acetyl-glucosamine (*O*-GlcNAcylation) that modulates various tumor-associated biological activities. By using a cutting-edge chemical proteomic methodology for intact glycopeptide analysis, we show herein that *O*-GlcNAcylation of Keratin 18 (K18) coordinates the tricarboxylic acid (TCA) cycle enzymes, namely isocitrate dehydrogenases (IDHs), to promote CCA progression. Mechanistically, site-specific *O*-GlcNAcylation of K18 on Ser 30 stabilizes K18, which benefits the expression of cell cycle checkpoints to enhance cell cycle progression and cell growth. Interaction with IDHs down-regulates the level of citrate and isocitrate, while up-regulates the level of α-ketoglutarate (α-KG). Our study thus expands the current understanding of protein *O*-GlcNAcylation, and adds another dimension of complexity to post-translational control over metabolism and tumorigenesis.

## Introduction

Cholangiocarcinoma (CCA), also known as bile duct cancer, constitutes a constellation of malignancies emerging in the biliary tree. CCA is the second most common primary hepatic malignancy after hepatocellular carcinoma (HCC), accounting for approximately 15% of all primary liver tumors, and its incidence is increasing worldwide.^1–3^ CCA is typically asymptomatic in the early stages and difficult to cure at late stages, which highly compromises therapeutic options and leads to a dismal prognosis.^4–7^ The discoveries of early diagnostic markers are, therefore, of great significance for improving CCA diagnosis and therapeutics.^4,8^

A growing appreciation for the physiological roles of glycans, either in glycoconjugates (*e.g.,* glycoproteins, glycolipids, proteoglycans) or in their free form, have resulted in increasing attention in their functional elucidation.^9–13^ Of note, β-*O*-linked *N*-acetylglucosamine (GlcNAc) is a dynamic glycosylation attached to serine and threonine of nucleocytoplasmic and mitochondrial proteins. This modification is dynamically concerted by a pair of enzymes, *O*-GlcNAc transferase (OGT) and *O*-GlcNAcase (OGA).^14^ *O*-GlcNAcylation has been identified in more than 5,000 human proteins and exhibits complex crosstalk with protein phosphorylation.^15,16^ Accumulating evidence has shown that *O*-GlcNAcylation appears to modulate various biological activities (*e.g.*, epigenetics, transcription, cellular metabolism) in response to environmental cues, and alternation in *O*-GlcNAcylation underlies neurodegenerative diseases, diabetes and various cancers.^17–19^ Aberrant *O*-GlcNAcylation actively contributes to pathophysiological steps of tumor progression, by regulating cancer cell signaling, proliferation, invasion, angiogenesis, metastasis, cell-matrix interaction and immune modulation both *in vitro* and *in vivo*.^12,20,21^ However, the mechanistic elucidation by which protein-specific *O*-GlcNAcylation contributes to CCA metabolic reprogramming and tumorigenesis remains largely elusive. Herein we present the systematic perception of *O*-GlcNAcylation, OGT and OGA in human CCA samples, and then perturbed the CCA cells with a metabolically-based intact glycopeptide analysis strategy, to profile global *O*-GlcNAcylation with glycan composition and glycosylation site resolution. We also provide mechanistic insight as to how Keratin 18 *O*-GlcNAcylation affects and choreographs the tricarboxylic acid (TCA) cycle in CCA progression.

## Results

### *O*-GlcNAcylation is up-regulated in CCA

We started by investigating the clinical relevance of *O*-GlcNAcylation in CCA. We first analyzed the *O*-GlcNAc, and the expression of OGT and OGA in 21 pairs of human resected tumor tissues and adjacent normal bile duct (BD) using immunohistochemistry (IHC) (**Figure 1A**). The levels of *O*-GlcNAc and OGT were distinctively elevated in tumor tissues compared with normal tissues, consistent with the IHC staining scores in these CCA patients (P < 0.001, **Figure 1B**). However, the level of OGA did not show distinctive changes between the two groups (**Figures 1A and 1B**). We then analyzed the *O*-GlcNAc expression levels in additional 15 peritumoral/tumor tissue pairs from CCA patients. Again, *O*-GlcNAcylation and OGT were identified to be up-regulated in tumor tissues (**Figures 1C and S1**), and quantitative analysis of the signals confirmed that the increases are statistically significant (P < 0.01 and 0.05, respectively by Student’s t-tests, two-tailed) (**Figure 1D**). Notably, mRNA expression of *OGT* and *OGA* was escalated from the Cancer Genome Atlas (TCGA) and four CCA disease datasets (GSE32879, GSE107943, GSE119336, GSE76297) (**Figures S2A and S2B**). CCA patients with high levels of *OGT* and low levels of *OGA* gene expression displayed worse overall survival (OS) according to the Kaplan-Meier survival analysis, indicating the pivotal role of *O*-GlcNAcylation in CCA (**Figures S2C and S2D**). We further assessed the *O*-GlcNAc, OGT and OGA expressions in three human CCA cell lines, namely HCCC-9810, RBE and HuCCT1, and the Human Intrahepatic Biliary Epithelial Cell (HIBEpiC) control cell line (**Figure 1E**). *O*-GlcNAcylation level increased in all CCA cells compared with HIBEpiC cells, while mRNA expression of *OGT* and *OGA*, as measured by quantitative real-time PCR (qRT-PCR), exhibited varied expression levels (**Figure 1F**). These data altogether pinpointed the up-regulation of *O*-GlcNAcylation as a common event in CCA samples.

**Figure 1.**
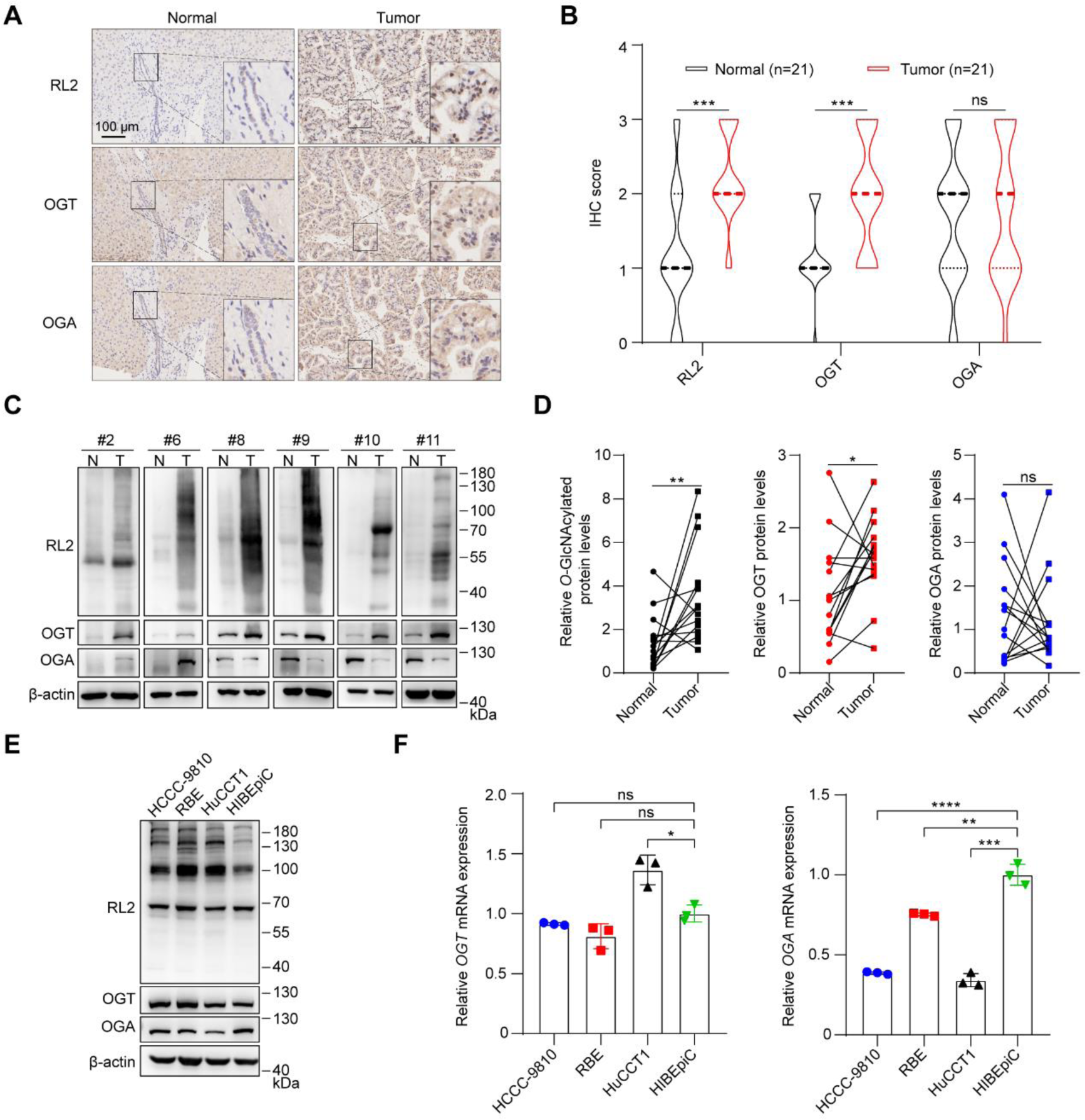
Dysregulation of *O*-GlcNAcylation in human CCA. **(A)** Representative images of RL2 (anti-*O*-GlcNAc), OGT and OGA staining by immunohistochemistry (IHC). Scale bar: 100 μm. **(B)** The analysis of IHC staining scores of RL2 (anti-*O*-GlcNAc), OGT and OGA in 21 pairs of CCA tumor tissues and adjacent normal bile ducts. Level of staining: 0-negative, 1-weakly positive, 2-positive, 3-strongly positive. **(C)** Representative images of *O*-GlcNAcylated protein, OGT and OGA levels from six CCA tumor tissues (T) and adjacent normal tissues (N) by Western blot analysis. Equal loading was confirmed using β-actin. **(D)** Densitometric analysis of *O*-GlcNAcylated protein, OGT and OGA levels from 15 pairs of CCA tumor tissues and adjacent normal tissues by Western blot analysis. **(E)** *O*-GlcNAcylated protein, OGT and OGA levels of three CCA cell lines and the Human Intrahepatic Biliary Epithelial Cell (HIBEpiC) control cell line by Western blot analysis. Equal loading was confirmed using β-actin. **(F)** The relative expression level of *OGT* or *OGA* mRNA in three CCA cells and HIBEpiC cells was determined by quantitative real-time PCR (qRT-PCR). CCA, cholangiocarcinoma; *O*-GlcNAc, β-*O*-linked *N*-acetylglucosamine; *O*-GlcNAcylation, *O*-linked β-*N*-acetylglucosaminylation; OGT, *O*-GlcNAc transferase; OGA, *O*-GlcNAcase. Data were shown as the mean ± standard deviation (SD); statistical significance was determined by Student’s t-tests (two-tailed, *P < 0.05, **P < 0.01, ***P < 0.001, ****P < 0.0001, ns, not significant).

### Global *O*-GlcNAcylation contributes to CCA cell proliferation

Previous reports by the Wongkham group indicated the significance of *O*-GlcNAcylation in controlling the metastatic ability of CCA cells via nuclear translocation of NF-κB and heterogeneous nuclear ribonucleoprotein-K (hnRNP-K).^22–24^ We therefore postulated that modulation of *O*-GlcNAc levels would alter CCA oncology phenotypes (*i.e.*, proliferation, apoptosis, cell cycle), by targeting pathways known to regulate CCA progression. To do so, we first used chemical tools to inhibit OGT and OGA. Ac_4_5SGlcNAc (designated as 5S thereof) is a known inhibitor of OGT that acts as a metabolic precursor to form UDP-5SGlcNAc, and OGA can be selectively and effectively inhibited by Thiamet-G (designated as TMG thereof) (**Figure 2A**).^25,26^ Treatment of all three CCA cell lines with 5S or TMG led to the robust decrease or increase of *O*-GlcNAc modification level on proteins, in a time- and dose-dependent manner (**Figures 2B, S3A, S4A, and S5**), in accordance with previous observations. At the same time, cellular viability was assessed using a cell counting kit-8 (CCK-8). The overall cytotoxicity in HuCCT1 cells was governed by the inhibition of OGT rather than OGA, suggesting that lowering *O*-GlcNAcylation will induce cell death (**Figure 2C**). Similar effects were also observed in RBE and HCCC-9810 cells in a dose-dependent manner (**Figures S3B and S4B**). Moreover, suppression of OGT promoted CCA cell apoptosis, as evidenced by the increased percentage of both early apoptosis (FITC^+^/PE^-^) and late apoptosis (FITC^+^/PE^+^) after OGT silencing upon 5S incubation (**Figures 2D and S3C**). No remarkable effects of TMG as an anti-apoptotic inhibitor were detected (**Figures 2D and S3C**). Mounting reports implied that *O*-GlcNAc plays a multifaceted role during the cell cycle, and incongruousness arises partly due to different physiological cues.^27^ We examined the cell cycle distribution of HuCCT1 and RBE cells using flow cytometric analysis after incubation with 5S or TMG at varied concentrations (**Figures 2E and S3D**). Interestingly, cell cycle distribution of HuCCT1 cells was majorly found in the G2/M phase, while RBE cells were arrested in the S phase when treated with 5S, but not under TMG-treated scenarios (**Figures 2E and S3D**). It is therefore sufficient to conclude that an appropriate *O*-GlcNAc level is critical for all cell cycle phases. Additionally, we blotted the classical biomarkers for cell cycle and apoptosis after perturbation of *O*-GlcNAcylation using 5S and TMG, respectively (**Figures 2F, S3E and 2J**). There were no notable differences in these biomarkers for the TMG-treated group, yet the 5S-treated group presented global up-regulation on apoptosis markers and down-regulation on the cell cycle checkpoints (**Figures 2F, S3E and 2J**). To further elucidate the correlation between *O*-GlcNAcylation and cancer progression, we complemented three independent small interfering RNAs (siRNAs) for *OGT* or *OGA* knockdown (**Figures 2G and S3F**), all of which showed potent knockdown efficacy with the corresponding *O*-GlcNAc processing enzymes (*i.e.*, OGT or OGA), meanwhile maintaining the desired catalytic activities (**Figures 2H and S3G**). Clonogenic assay both qualitatively and quantitatively validated that knockdown of *OGT* in HuCCT1 and RBE cells clearly reduced cell proliferation and presented an anti-tumor effect (**Figures 2I and S3H**). These biological characterizations reciprocally integrate with chemical tools, emphasizing that global *O*-GlcNAcylation may promote the development and progression of cholangiocarcinoma.

**Figure 2.**
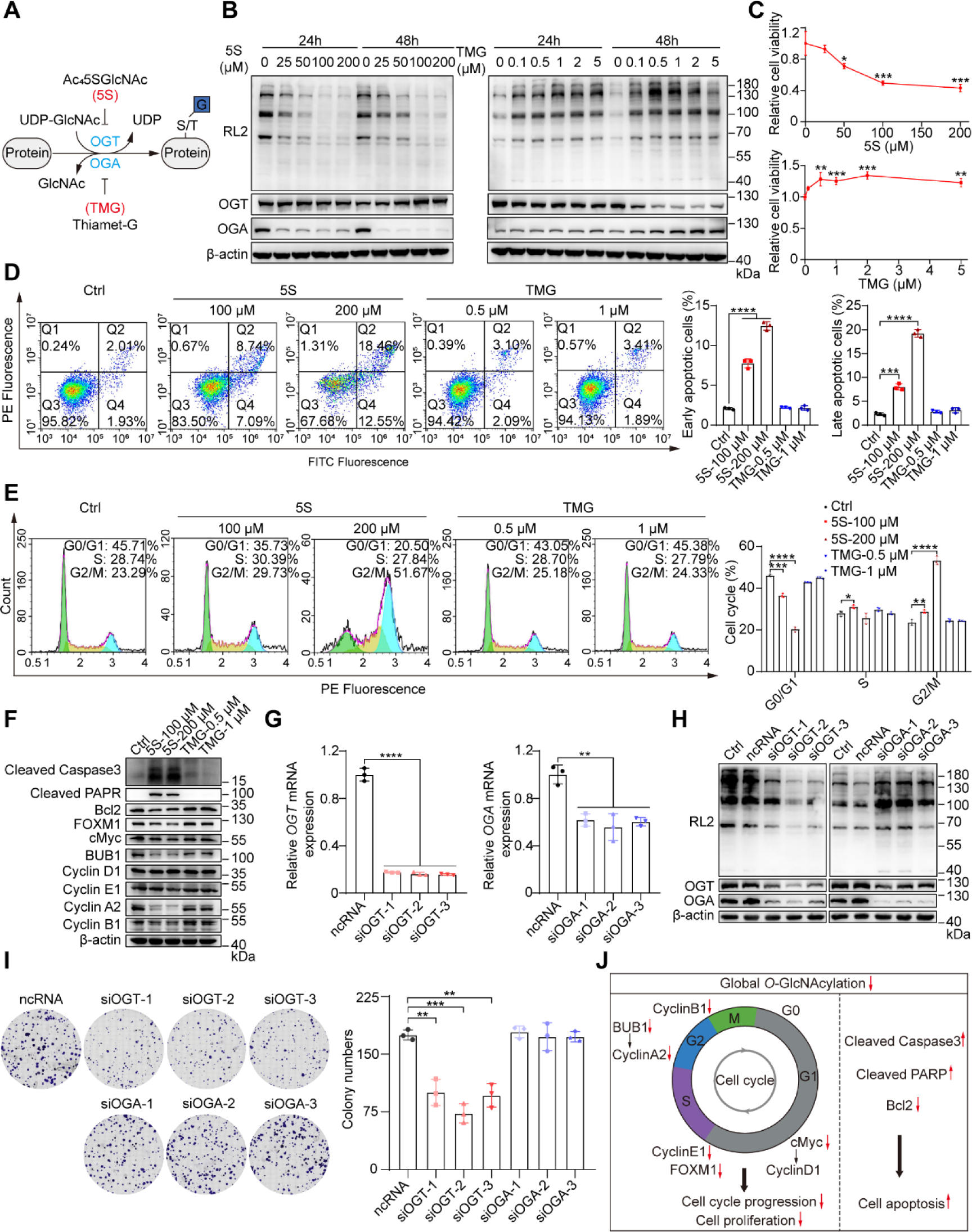
*O*-GlcNAcylation promotes CCA cell proliferation. **(A)** Schematic of protein *O*-GlcNAcylation processes and inhibition. A pair of enzymes, OGT and OGA, catalyze the addition/removal of single *O*-GlcNAc, respectively. Ac_4_5SGlcNAc (5S) is an inhibitor of OGT, and Thiamet-G (TMG) is an inhibitor of OGA. **(B)** Time and dose-dependent analysis of the *O*-GlcNAcylated protein, OGT and OGA levels in 5S- or TMG-treated HuCCT1 cells by Western blot. Equal loading was confirmed using β-actin. **(C)** Cytotoxicity assay of 5S- or TMG-treated HuCCT1 cells. The cells were treated with 5S or TMG at various concentrations for 48 h. **(D)** Cell apoptosis assay of 5S- or TMG-treated HuCCT1 cells. The bivariate density plot in flow cytometry indicated the cell population of early apoptotic cells (FITC^+^/PE^-^) and late apoptotic cells (FITC^+^/PE^+^). Quantitative analysis was shown in the right panel. **(E)** Cell cycle distribution assay of 5S- or TMG-treated HuCCT1 cells. Histogram plot in flow cytometry indicated the percentage of cell populations in the G0/G1, S, or G2/M phase. Quantitative analysis was shown in the right panel. **(F)** Cell cycle and apoptosis marker analysis of 5S- or TMG-treated HuCCT1 cells by Western blot. Protein levels of cleaved Caspase3, cleaved PARP (apoptotic markers); Bcl2 (anti-apoptotic marker); cMyc, Cyclin D1, FOXM1, Cyclin E1(G1/S transition markers); BUB1, Cyclin A2, Cyclin B1 (G2/M transition markers) were analyzed. Equal loading was confirmed using β-actin. **(G)** The *OGT* or *OGA* mRNA expression of small interfering RNA (siRNA)-treated HuCCT1 cells by qRT-PCR. Random RNA was used as the negative control (ncRNA). Three independent siRNAs targeting *OGT* (siOGT1-3) or *OGA* (siOGA1-3) were shown. **(H)** Silencing efficacy of siOGT or siOGA in HuCCT1 cells. The *O*-GlcNAcylated protein, OGT and OGA levels were analyzed by Western blot. Equal loading was confirmed using β-actin. **(I)** Clonogenic assay of HuCCT1 cells transfected with ncRNA, siOGTs or siOGAs. Colony numbers were quantified in the right panel. **(J)** Putative schematic model of *O*-GlcNAcylation regulation in CCA progression. Data were shown as the mean ± SD; statistical significance was determined by Student’s t-tests (two-tailed, *P < 0.05, **P < 0.01, ***P < 0.001, ****P < 0.0001).

### Chemical enrichment and profiling of intact *O*-GlcNAcylated glycopeptides in CCA

Since imbalanced *O*-GlcNAcylation impacts the process of cancer progression in CCA, we next explored to systematically enrich, identify and profile intact *O*-GlcNAcylated glycopeptides with matched information on glycosylation sites and glycan compositions. We adopted Click-iG, a comprehensive platform that amalgamates metabolic oligosaccharide engineering (MOE) of selected, clickable unnatural sugar probes for *O*-GlcNAcylated protein enrichment, and a customized pGlyco3 search engine for intact glycopeptide annotation.^28–30^ In brief, azidosugars were metabolically incorporated into various glycans. The azido-containing glycoproteins were reacted with a three-module alkyne-photocleavable linker-biotin tag (alkyne-PC-biotin), digested with trypsin, and enriched using streptavidin beads. After photocleavage with 365 nm ultraviolet (UV), the released click-labeled glycopeptides were subjected to liquid chromatography-tandem mass spectrometry (LC-MS/MS) analysis, with preferred glycopeptide fragmentation strategies, namely, stepped collision energy based higher-energy collisional dissociation (sceHCD) followed by product-dependent electron transfer/higher-energy dissociation (sceHCD-pd-EThcD) (**Figure 3A**). 1,6-di-*O*-propionyl-*N*-azidoacetylgalactosamine (1,6-Pr_2_GalNAz), an optimized monosaccharide *O*-GlcNAc chemical reporter with minimal nonspecific S-glyco-modification and cytotoxicity, was employed in the experiment (**Figure S6**).^31^ We first evaluated the metabolic efficacy of 1,6-Pr_2_GalNAz in the CCA and HIBEpiC cell lines and observed azidosugar incorporation in a dose- and time-dependent manner (**Figures 3B and S7A**). Significant fluorescence labeling at nucleocytoplasmic regions was achieved when 1,6-Pr_2_GalNAz-treated cells were permeabilized and reacted with alkyne-AZDye-488, via the ligand-assisted copper (I)-catalyzed azide-alkyne cycloaddition (CuAAC), in agreement with the localization of *O*-GlcNAc glycosylation (**Figures 3C and S7B**). Similar quantitative data on all four cell lines were observed by flow cytometry (**Figure 3D**). In view of these results, HuCCT1 and HIBEpiC cells incubated with 200 μM 1,6-Pr_2_GalNAz for 48 h were used as the standardized condition for glycoproteomic analysis.

**Figure 3.**
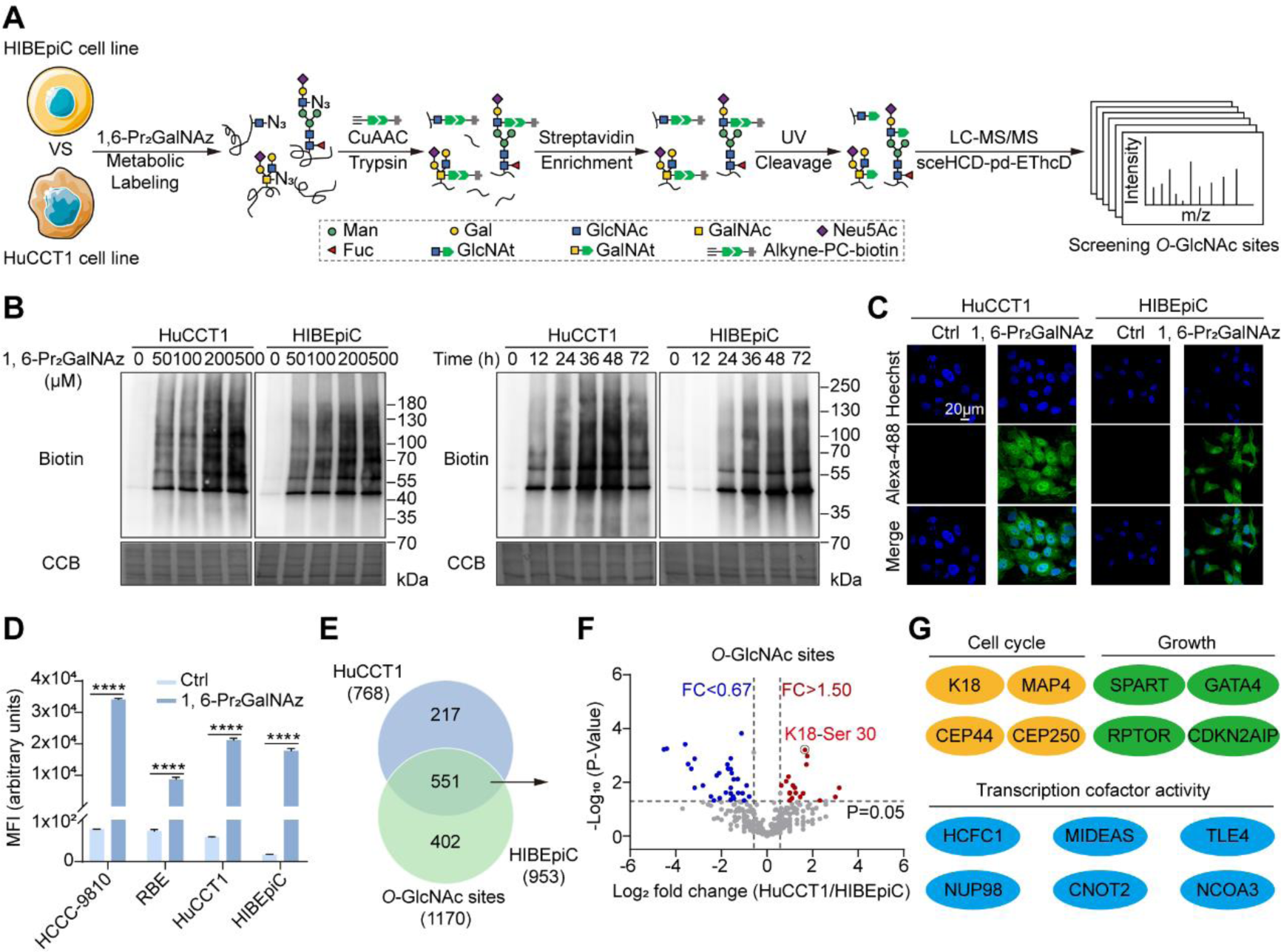
Profiling of protein *O*-GlcNAcylation in CCA by Click-iG. **(A)** Schematic of the Click-iG strategy in CCA analysis. HuCCT1 and HiBEpiC cells are metabolically incorporated with 1,6-di-*O*-propionyl-*N*-azidoacetylgalactosamine (1,6-Pr_2_GalNAz), reacted with alkyne-PC-biotin via click chemistry, digested and enriched by streptavidin beads, followed by photocleavage release of glycopeptides for whole glycopeptide analysis. **(B)** Western blot analysis of HuCCT1 and HiBEpiC cells treated with 1,6-Pr_2_GalNAz at various concentrations for different times. The cell lysates were reacted with alkyne-biotin via copper(I)-catalyzed azide-alkyne cycloaddition (CuAAC) and blotted using anti-biotin. Equal loading was confirmed using Coomassie brilliant blue staining (CBB). **(C)** Confocal fluorescence imaging of HuCCT1 and HIBEpiC cells treated with 1,6-Pr_2_GalNAz at 0 or 200 μM for 48 h. The cells were washed, labeled with alkyne-AZDye-488, and analyzed. Scale bar: 20 μm. **(D)** Flow cytometry analysis of three CCA cells and HIBEpiC cells treated with 1,6-Pr_2_GalNAz at 0 or 200 μM for 48 h. The cells were washed, reacted with alkyne-biotin and Alexa Flour 488-streptavidin, and analyzed. **(E)** Total numbers of *O*-GlcNAc sites identified in HuCCT1 and HIBEpiC cells in three independent experiment runs. **(F)** Volcano plots showing the average log_2_ fold change (HuCCT1/HIBEpiC) for *O*-GcNAc sites quantified in three biological replicates and P-values. *O*-GcNAc sites with P-value < 0.05 and a fold change > 1.50 (red) or < 0.67 (blue) were considered as up-regulated or down-regulated *O*-GcNAc sites, respectively. **(G)** *O*-GlcNAcylated proteins involved in the cell cycle, growth and transcription cofactor activity. sceHCD-pd-EThcD, stepped collision energy based higher-energy collisional dissociation followed by product-dependent electron transfer/higher-energy dissociation; Man, mannose; Gal, galactose; Glu, glucose; GlcNAc, *N*-acetylglucosamine; GalNAc, *N*-acetylgalactosamine; Neu5Ac, *N*-acetylneuraminic acid; Fuc, fucose; GlcNAt, *N*-(4-aminomethyl)-triazolylacetylglucosamine; GalNAt, *N*-(4-aminomethyl)-triazolylacetylgalactosamine; alkyne-PC-biotin, alkyne-photocleavable linker-biotin tag. Data were shown as the mean ± SD; statistical significance was determined by Student’s t-tests (two-tailed, ****P < 0.0001).

By using the established streamlined MS data analysis and annotation procedure (**Figure S8**), we then mapped a total of 1,170 *O*-GlcNAc sites on 368 *O*-GlcNAcylated proteins from three replicate experiments using the Click-iG workflow (**Figures 3A, 3E, S9A and S10A)**. In combination, the Click-iG strategy yielded pan-scale intact glycopeptide information (*i.e.*, intact glycosites, glycosites, glycoproteins) on HuCCT1 and HIBEpiC cells (**Figure S10B**). For instance, a total of 1,872 intact glycosites were identified in HuCCT1 cells, of which 1,094 overlapped with those identified in HIBEpiC cells, indicating the cell-type specific feature of protein glycosylation (**Figure S10B, left panel**). By glycan classification, we identified a total of 3,164 intact glycosites, which also consisted of 1,421 intact *N*-linked glycosites and 573 intact mucin-type *O*-linked glycosites (**Figure S10C**). Sequence visualized using a probabilistic approach around the identified glycosites revealed the typical sequon/motif for *O*-GlcNAcylation, mucin-type *O*-linked glycosylation and *N*-linked glycosylation (**Figures S9B and S10D**).^32^ Remarkably, the Click-iG also provided in-depth glycan type and composition information from intact glycosites (**Figures S11A, S11B, S12A, and S12B**). Gene Ontology (GO) analysis showed that the identified *O*-GlcNAcylated proteins concentrated in the nucleocytoplasmic region (**Figure S9C**). In addition, proteins carrying 23 up-regulated (fold change > 1.50, P < 0.05) and 36 down-regulated (fold change < 0.67, P < 0.05) *O*-GlcNAc sites in HuCCT1 cells were potently enriched using Click-iG (**Figure 3F; Table S2 and S3**), of which many protein regulators involved in the cell cycle and growth, as well as transcriptional processes, were identified to be *O*-GlcNAcylated (**Figures 3G and S13**). These results, collectively, demonstrate that Click-iG enables simultaneous and comprehensive profiling of *O*-GlcNAcylation (and other types of glycosylation) in the CCA and HIBEpiC cell lines, with intact glycosite level resolution.

### Keratin 18 is mainly *O*-GlcNAcylated at Ser 30

Keratins play a pivotal role in differentiation and tissue specialization and are highly regulated among various epithelia in a cell-specific manner.^33,34^ We noticed that Keratin 18 (K18) is on the list of up-regulated *O*-GlcNAcylated proteins in HuCCT1 cells compared to HIBEpiC control cells (**Figure 3F; Tables S2 and S3**). Keratin 18 plays various roles in intracellular scaffolding and cellular processes and is highly associated with malignant phenotype in digestive epithelia.^35–40^ Post-translational modification (PTM) including sumoylation,^41^ acetylation/methylation,^42^ *O*-GlcNAcylation^43^ and its reciprocal cross-talk with phosphorylation^44,45^ on K18 have been reported in previous works. To confirm the *O*-GlcNAcylation of K18, we incubated HuCCT1 and HIBEpiC cells with 1,6-Pr_2_GalNAz for 48 h, reacted with alkyne-biotin using CuAAC, and conducted pull-down procedures with streptavidin beads. Immunoblotting with anti-K18 demonstrated that K18 was modified with azidosugars (**Figure 4A, left panel**). To further verify the labeling results, we treated both cell lysates with a permissive β-1,4-galactosyltransferase mutant (Y289L GalT1), which transfers *N*-azidoacetylgalactosamine (GalNAz) from its uridine diphosphate activated precursor (UDP-GalNAz) to *O*-GlcNAc residues.^46,47^ Subsequent chemoselective click reaction with alkyne-biotin followed by streptavidin enrichment proved *O*-GlcNAcylation occurred on endogenous K18 (**Figure 4A, right panel**). The extent of *O*-GlcNAc modification on K18 in HuCCT1 cells was semi-quantitatively measured by labeling azides with alkynylated polyethylene glycol 5000 (alkyne-PEG_5KD_) in a mass shift assay. The stoichiometric ratio for *O*-GlcNAcylation, whether in *O*-β-*N*-azidoacetylglucosamine (*O*-β-GlcNAz) or in *O*-β-GlcNAc-β-1,4-GalNAz form, was approximately 50% (**Figure 4B**). Based on Click-iG, we mapped four *O*-GlcNAc modification sites (Ser 15, Ser 18, Ser 30 and Ser 31) on K18, of which Ser 30 and Ser 31 were previously reported. Interestingly, *O*-GlcNAcylation on Ser 49 was previously reported yet not found in our experiment setting.^43^ Of note, all four sites identified are well conserved in *Homo sapiens*, *Mus musculus* and *Rattus norvegicus*. (**Figures S14A and S14B**). To further determine the relative abundance of *O*-GlcNAc modification on these sites, the FLAG-tagged K18 mutants, K18^S15A^, K18^S18A^, K18^S15/18A^, K18^S30A^, K18^S31A^, K18^S30/31A^ and K18^S15/18/30/31A^ (K18^4A^), were transfected into HEK293T cells. The incorporated azides, either via MOE with 1,6-Pr_2_GalNAz, or Y289L GalT1 chemoenzymatic labeling, were reacted with alkyne-biotin for streptavidin capture and immunoblot analysis. Ser 30 appears as the major *O*-GlcNAcylation site in K18 (**Figures 4C and 4D**), in line with our chemical proteomic analysis results (**Figure 3F**). Alternatively, we pulled down with anti-FLAG beads for K18 enrichment, and then measured the *O*-GlcNAc level for FLAG-K18^WT^ or FLAG-K18^S30A^, using biotinylated signals introduced via the above-mentioned glyco-analytical methods. A significant loss of *O*-GlcNAcylation for FLAG-K18^S30A^ compared with FLAG-K18^WT^ was observed (**Figures 4E and 4F**). In addition, sceHCD-pd-EThcD based LC-MS/MS analysis annotated the K18 peptide with amino acid 28-45 (PVSSAASVYAGAGGSGSR) as an *O*-GlcNAcylated peptide at Ser 30 (**Figure 4G**).

**Figure 4.**
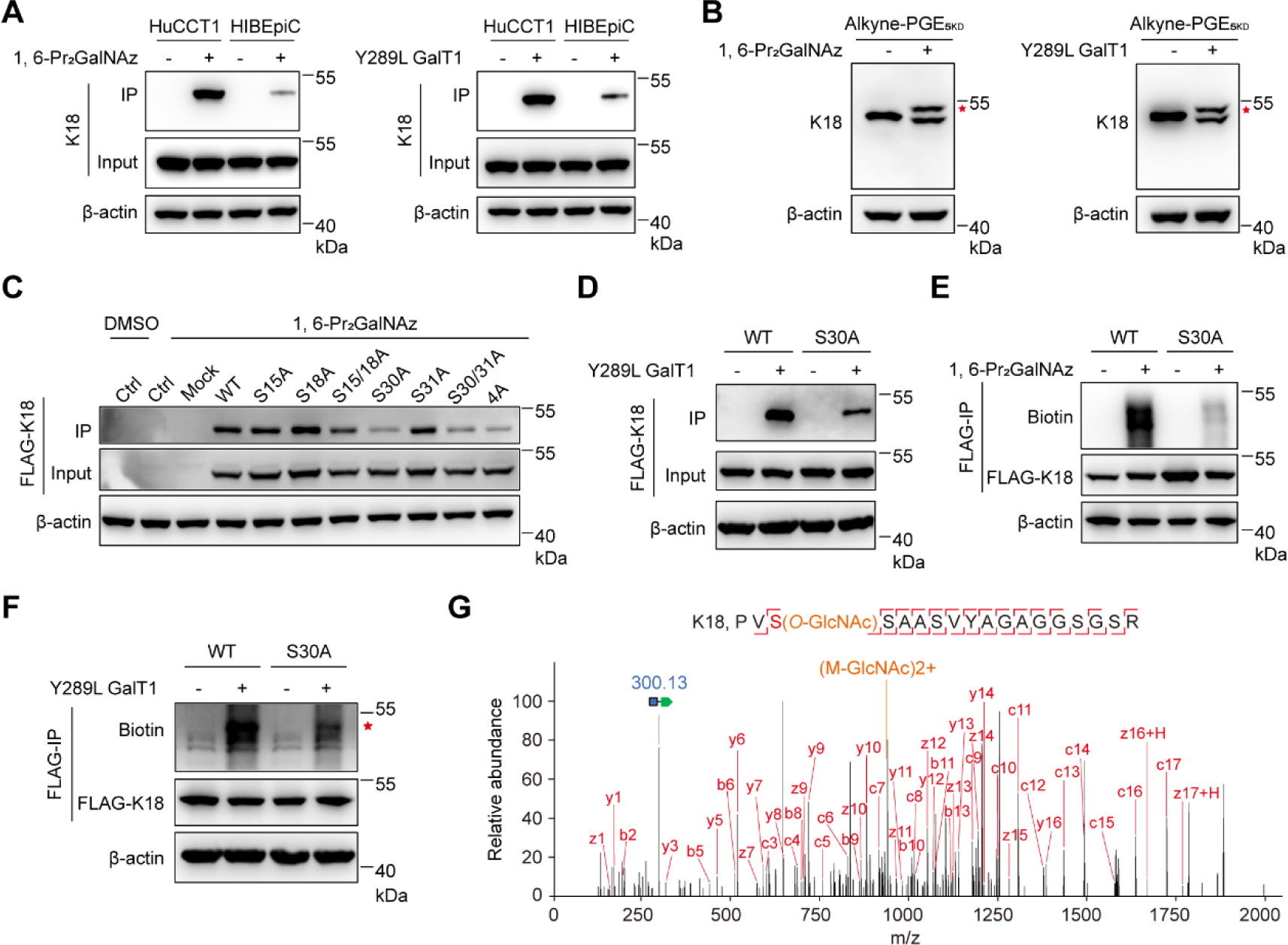
Keratin 18 is mainly *O*-GlcNAcylated at Ser 30. **(A)** Western blot analysis showing the Keratin 18 (K18) *O*-GlcNAcylation in HuCCT1 and HIBEpiC cells. The cells were incubated with 1,6-Pr_2_GalNAz, lysed, reacted with alkyne-biotin, and captured by streptavidin beads (left), or incubated with Y289L GalT1 and UDP-GalNAz in cell lysates, reacted with alkyne-biotin, and captured by streptavidin beads (right). **(B)** Western blot analysis showing the *O*-GlcNAcylation stoichiometry of K18 in HuCCT1 cells. The cells were incubated with 1,6-Pr_2_GalNAz, lysed, reacted with alkyne-PEG_5KD_ (left), or incubated with Y289L GalT1 and UDP-GalNAz in cell lysates, reacted with alkyne-PEG_5KD_ (right). The red asterisk indicated tagged O-GlcNAcylated K18. **(C)** Immunoblots analysis of K18 *O*-GlcNAcylation showing HEK293T cell lysates transfected with FLAG-tagged K18 (FLAG-K18) with wild type, single, double or quadruple mutations incubated with 1,6-Pr_2_GalNAz, lysed, and immunoprecipitated with streptavidin beads. **(D)** Immunoblot analysis showing HEK293T cells overexpressing FLAG-K18^WT^ or FLAG-K18^S30A^ incubated with Y289L GalT1 and UDP-GalNAz, and immunoprecipitated with streptavidin beads. **(E)** Immunoblot analysis showing the HEK293T cells overexpressing FLAG-K18^WT^ or FLAG-K18^S30A^ incubated with 1,6-Pr_2_GalNAz, lysed, reacted with alkyne-biotin, immunoprecipitated with anti-FLAG beads, and blotted with anti-biotin. **(F)** Immunoblot analysis showing the HEK293T cells overexpressing FLAG-K18^WT^ or FLAG-K18^S30A^ incubated with Y289L GalT1 and UDP-GalNAz, and immunoprecipitated with anti-FLAG beads, and blotted with anti-biotin. The red asterisk indicated tagged O-GlcNAcylated K18. **(G)** Representative MS2 spectrum of an *O*-GlcNAcylated peptide from K18 located on Ser 30. The matched fragment ions (red), diagnostic fragment ion (orange) and the GlcNAt fragment ion (blue) are labeled. Equal loadings were confirmed using β-actin in all Western blot analyses. UDP-GalNAz, UDP-*N*-azidoacetylglucosamine; Y289L GalT1, β-1,4-galactosyltransferase mutant; alkyne-PEG_5KD_, alkynylated polyethylene glycol 5000.

### *O*-GlcNAcylation of K18 promotes CCA proliferation and progression *in vitro* and *in vivo*

With the detailed glycosylation information for K18 at hand, we next asked whether K18 *O*-GlcNAc modification would impact cholangiocarcinoma phenotype(s). We first examined the K18 *O*-GlcNAcylation levels among typical human CCA cell lines and the normal HIBEpiC cell line. Elevated K18 *O*-GlcNAcylation was observed in all three CCA cell lines but not in HIBEpiC cells (**Figure 5A**). To scrutinize the importance of Ser 30 *O*-GlcNAcylation, we generated stable CCA cell lines with three independent targeting-resistant short hairpin RNA (shRNA) for K18 knockdown (**Figures S15A, S15B, S16A and S17A**), and then recovered K18 expression using either FLAG-K18-WT or FLAG-K18-S30A (**Figures 5B, S15C, S16B and S17B**). A systematic evaluation of *KRT18* mRNA expression and knockdown efficiency led to shK18-2 as the optimal construct (designated as shK18 thereof). CCK-8 and colony formation assays indicated that depletion of K18 gravely inhibited cell proliferation in HuCCT1 cells, which was rescued by re-expression of FLAG-K18-WT rather than FLAG-K18-S30A (**Figures 5C and 5D**). Similar results were also observed in RBE and HCCC-9810 cells (**Figures S16C, S16D, S17C and S17D**). These data imply that Ser 30 *O*-GlcNAcylation is functionally crucial in CCA proliferation *in vitro*. We next assessed the cell cycle distribution for each rescue cell line and found that FLAG-K18-S30A-rescued HuCCT1, RBE and HCCC-9810 cells showed an increase in S and G2/M phase arrest, partially in conformity with a previous report that *O-*GlcNAc on K18 is known to increase upon G2/M phase arrest (**Figures 5E, S16E and S17E**).^48^ Furthermore, immunoblot analysis on cell cycle biomarkers for these rescued cell lines displayed a similar correlation (**Figures 5F, S16F and S17F**).

**Figure 5.**
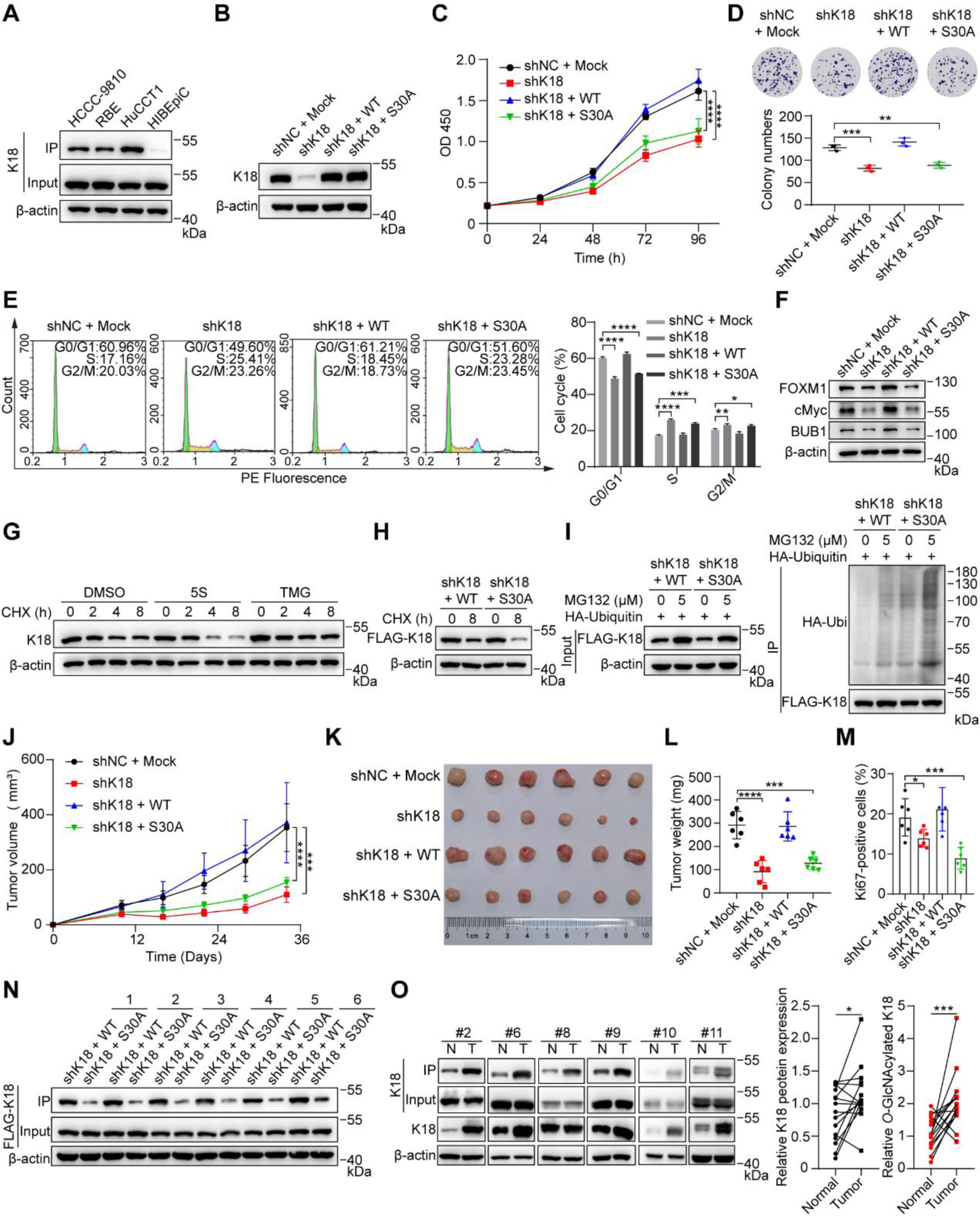
*O*-GlcNAcylation of K18 promotes CCA cell growth *in vitro* and *in vivo*. **(A)** Western blot analysis of K18 *O*-GlcNAcylation in three CCA cells and HIBEpiC cells. The cell lysates were incubated with Y289L GalT1 and UDP-GalNAz, reacted with alkyne-biotin, and immunoprecipitated by streptavidin beads. **(B)** Western blot analysis of K18 in HuCCT1 stable cell lines with small hairpin RNA K18 knockdown (shK18) and re-expression of shK18-resistant FLAG-K18 wild-type (shK18 + WT) or FLAG-K18 S30A (shK18 + S30A). Random small hairpin RNA with an empty vector (shNC + Mock) was used as a negative control. **(C)** CCK-8 analysis of HuCCT1 stable cell lines. Optical Density (OD) 450 was measured for cell viability. **(D)** Clonogenic assay of cell proliferation in HuCCT1 stable cell lines. Colony numbers were quantitatively analyzed at the bottom. **(E)** Cell cycle distribution assays of HuCCT1 stable cell lines. Histogram plot in flow cytometry indicated the percentage of cell populations in the G0/G1, S, or G2/M phase. Quantitative analysis was shown in the right panel. **(F)** Cell cycle marker analysis of HuCCT1 stable cell lines by Western blot. Protein levels of FOXM1, cMyc (G1/S transition markers); BUB1(G2/M transition marker) were analyzed. **(G)** Degradation analysis of K18 in HuCCT1 cells by Western blot. The cells were incubated with DMSO (vehicle), 200 μM 5S or 1 μM TMG for 48 h, followed by treatment with 10 μg/ml cycloheximide (CHX) for up to 8 h. **(H)** Degradation analysis of K18 in HuCCT1 shK18 + WT or shK18 + S30A stable cell lines by Western blot. The cells were incubated with DMSO (vehicle) or 10 μg/ml CHX for 8 h. **(I)** Ubiquitination analysis of K18 in HuCCT1 shK18 + WT or shK18 + S30A stable cell lines. The cells were transfected with HA-ubiquitin, incubated with 5 μM MG-132 (proteasome inhibitor) for 20 h, lysed and captured with anti-FLAG beads (left). Anti-HA blot demonstrated the ubiquitination of immunoprecipitated FLAG-K18 (right). **(J-N)** A tumorigenesis assay was performed by subcutaneous injection of HuCCT1 cells with shNC + Mock, shK18, shK18 + WT and shK18 + S30A into the right flanks of nude mice (n = 6). **(J)** Tumors generated by xenograft HuCCT1 stable cells were measured every six days from day 10. After 34 days, tumors were dissected, **(K)** photographed and **(L)** weighed. **(M)** Ki67 (a marker of cell proliferation) positive area analysis of xenograft tumors through immunohistochemistry. Scale bar: 100 μm. **(N)** Western blot analysis of K18 *O*-GlcNAcylation in xenograft tumors generated from stable HuCCT1 cell lines. **(O)** Representative images and densitometric analysis of K18 and its *O*-GlcNAcylation levels from 15 pairs of CCA tumor tissues (T) and adjacent normal tissues (N) by Western blot analysis. Equal loadings were confirmed using β-actin in all Western blot analyses. CCK-8, cell counting kit-8. Data were shown as the mean ± SD; statistical significance was determined by Student’s t-tests (two-tailed, *P < 0.05, **P < 0.01, ***P < 0.001, ****P < 0.0001).

Given that *O*-GlcNAcylation on K18 was reported to regulate its solubility, filament organization, and stability,^49^ we tested whether the stability of K18 is modulated by *O*-GlcNAcylation at Ser 30. Previous reports from the Rajiv lab exhibited that the half-life for K18 is regulated by *O*-GlcNAcylation in human hepatocytes (Chang) cell line.^44,49^ The CCA cell lines were treated with cycloheximide (CHX) to block protein synthesis for an immunoblot chase assay. OGT inhibition by 5S considerably accelerated the K18 degradation, while silencing of OGA by TMG poses minimal effect on its rate of decay (**Figures 5G, S16G and S17G**). Likewise, FLAG-K18-WT was more stable than FLAG-K18-S30A after 8 h of protein lifespan (**Figures 5H, S16H and S17H**). Correlatively, an increased level of ubiquitination was observed in FLAG-K18-S30A (**Figures 5I, S16I and S17I**). These findings indicate that K18 stability was enhanced through up-regulation of its *O*-GlcNAcylation at Ser 30, as well as the inhibition of the corresponding ubiquitination.

To decipher the effect of K18 *O*-GlcNAcylation on tumor growth *in vivo*, we injected BALB/c nude mice with shRNA negative control with an empty vector (shNC + Mock), shK18, shK18 + WT, shK18 + S30A stable HuCCT1 cell lines, and quantitatively measured tumor formation. K18 depletion and its mutant at Ser 30 greatly repressed tumor growth rate, tumor size/weight, and the Ki67 positive percentage (a marker to determine cancer cell proliferation) (**Figures 5J-M, S18A, and S18B**). In parallel, tumor tissues dissected from HuCCT1 cells expressing FLAG-K18-WT showed a higher level of K18 *O*-GlcNAcylation than the corresponding cells expressing FLAG-K18-S30A (**Figure 5N**). Luckily, we observed that both the protein expression level and the *O*-GlcNAcylation level for K18 were significantly enhanced when advancing toward the clinical CCA tumor tissues compared with adjacent normal tissues (**Figures 5O and S19**). These results are able to recapitulate most of the effects observed *in vitro* and solidify the hypothesis that *O*-GlcNAcylation of K18 promotes CCA progression *in vivo*.

### K18 *O*-GlcNAcylation promotes K18-Isocitrate dehydrogenase interaction to regulate the TCA cycle in CCA

*O*-GlcNAcylation serves as a nutrient rheostat in a myriad of physiological contexts, especially in metabolically active organs such as the liver. The tight interconnection between *O*-GlcNAcylation and glucose metabolism prompted us to explore the mechanistic insight as to how K18 *O*-GlcNAcylation affects its interaction.^50–52^ We transfected HuCCT1 shK18 stable cell line with FLAG-K18^WT^ or FLAG-K18^S30A^ with equivalent protein expression level, co-immunoprecipitated the K18-interacting proteins with anti-FLAG beads and subjected to LC-MS/MS analysis (**Figures S20A and S20B**). We identified 858 up-regulated (fold change > 1.50, P < 0.05) interacting proteins in FLAG-K18^WT^ HuCCT1 cells compared to FLAG-K18^S30A^ cells (**Figure 6A**), the majority of which were closely related to the TCA cycle shown by the Kyoto Encyclopedia of Genes and Genomes (KEGG) pathway analysis (**Figure 6B**). To further elucidate the K18 binding partners in the TCA cycle, we analyzed 8 out of 13 up-regulated TCA enzymes, by immunoblotting with their corresponding antibodies (**Table S4**). We noticed an apparent signal decrease in isocitrate dehydrogenase (IDH) families, including IDH2, IDH3A, IDH3B and IDH3G, when *O*-GlcNAc modification at Ser 30 of K18 was functionally amputated (**Figure 6C**). The IDH enzymes are responsible for the oxidative decarboxylation of isocitrate. To further corroborate these results, we performed relative quantification of cellular metabolites using LC-MS/MS analysis. We found that in the absence of *O*-GlcNAcylation at Ser 30 on K18, the rescued cells accumulated higher levels of citrate, aconitate and isocitrate, but lower levels of pyruvate, α-ketoglutarate, succinate, fumarate and malate, compared to K18 wild type rescued cells (**Figure 6D**). These data suggest a strong correlation between K18 *O*-GlcNAcylation and IDH catalytic activities. Similar results were observed in RBE-rescued cells (**Figures S21A and S21B**). *In toto*, we conclude that in cholangiocarcinoma, up-regulated OGT level will inevitably enhance the K18 *O*-GlcNAcylation, which increases its interaction with isocitrate dehydrogenases in the TCA cycle, one of the major metabolic pathways to regulate glucose metabolism, to coordinate the cell proliferation and CCA progression (**Figure 6E**).

**Figure 6.**
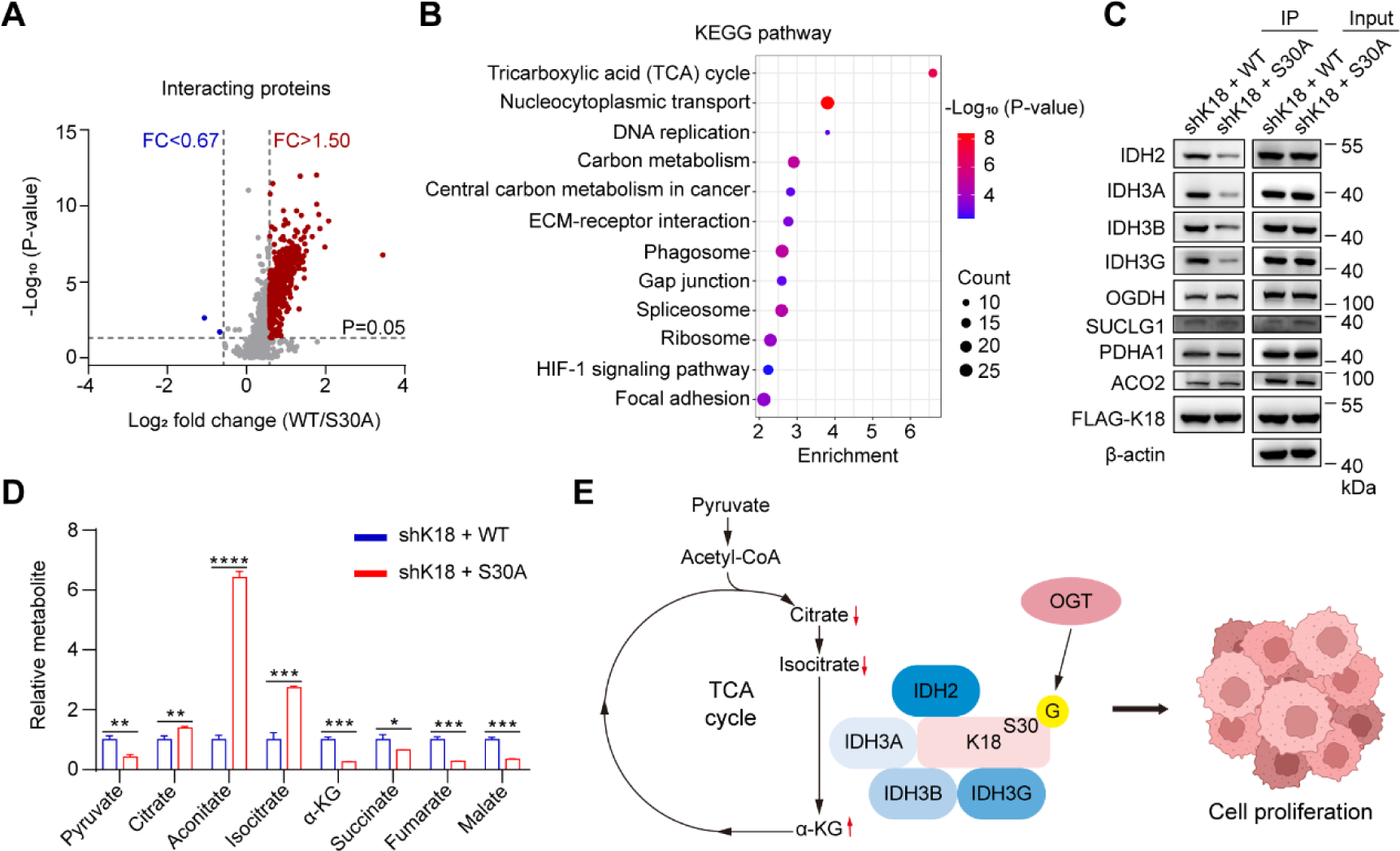
*O*-GlcNAcylation of K18 regulates the metabolism of TCA cycle. **(B)** Volcano plot showing the relative abundance of interacting proteins with K18 in HuCCT1 shK18 stable cells transfected with FLAG-K18^WT^ or FLAG-K18^S30A^. the average log_2_ flod change for proteins quantified in three biological replicates and P-values. Proteins with P-value < 0.05 and a fold change > 1.50 (red) or < 0.67 (blue) were considered as up-regulated or down-regulated interacting proteins, respectively. **(C)** KEGG enrichment analysis for the upregulated interacting proteins with K18 in HuCCT1 shK18 cells transfected with FLAG-K18^WT^ compared with FLAG-K18^S30A^. **(D)** The analysis of enzymes in the TCA cycle interacting with K18 of HuCCT1 stable cell lines (shK18 + WT and shK18 + S30A). Protein levels of IDH2, IDH3A, IDH3B, IDH3G, OGDH, SUCLG1, PDHA1, ACO2 and FLAG-K18 were analyzed by Western blot and immunoprecipitation analysis. Equal loadings were confirmed using β-actin. **(E)** Relative abundance of metabolites of the TCA cycle in HuCCT1 stable cell lines (shK18 + WT and shK18 + S30A). **(F)** Proposed functional action of K18 *O*-GlcNAcylation in promoting CCA progression. KEGG, Kyoto Encyclopedia of Genes and Genomes; α-KG, α-ketoglutaric acid; TCA, tricarboxylic acid. Data were shown as the mean ± SD; statistical significance was determined by Student’s t-tests (two-tailed, *P < 0.05, **P < 0.01, ***P < 0.001, ****P < 0.0001).

## Discussion

The glycosylation landscape of CCA, and how it affects CCA diagnosis and prognosis, has recently emerged as an appealing research field.^53,54^ Glyco-biomarkers such as CA19-9 and CEA are up-regulated in CCA and exhibit diagnostic value.^55,56^ Glyco-analytical methodologies including lectin blot,^57–59^ glycan microarray^60^ and glycoproteomic analysis^61–64^ have been utilized to probe the glycobiology in CCA. In addition, abnormal sialylation,^65–68^ fucosylation^69,70^ and *O*-linked *N*-acetylgalactosamine (GalNAc) modification (*O*-GalNAcylation)^60,71–73^ have also been implicated in prognostic outcome in human cholangiocarcinoma. Changes in *O*-GlcNAc levels and the cycling enzyme pair OGT/OGA in a myriad of tumor types have also been reviewed.^54,74^ However, a thorough investigation on how *O*-GlcNAcylation affects CCA progression, based on chemical glycoproteomic analysis has never been previously reported, to the best of our knowledge. In this study, we utilized a chemical glycoproteomic strategy Click-iG, to mechanistically investigate the molecular basis of *O*-GlcNAcylation in CCA, as exemplified by K18 *O*-GlcNAcylation at Ser 30. Alternative CCA glyco-biomarker candidates are also included within the intact glycopeptide analysis data (**Tables S2 and S3**) that await validation. It is expected to enhance our current knowledge of CCA progression and may reveal novel targets for this highly lethal malignancy.

We conduct the systematic analysis of *O*-GlcNAcylation, OGT and OGA in cholangiocarcinoma from clinical samples, and demonstrate a universal increase in *O*-GlcNAcylated proteins, in correlation with OGT up-regulation. Contrarily, we do not observe a down-regulation of OGA protein level in CCA as previously reported,^22^ further illustrating the individual variability and complexity of clinical samples. Hence, we choose the CCA and HIBEpiC cell lines, which offer an easy-to-manipulate, cost-effective and stable platform. The roles of *O*-GlcNAcylation on metastasis, migration and invasion abilities in CCA cells were previously emphasized.^23,24,75^ Thus, we focus on the biological influence of *O*-GlcNAcylation on cell apoptosis, cell cycle and proliferation in the CCA cells. OGT and OGA are essential in mammals, the mutation in *OGT* and/or *OGA* requires conditional alleles, and studies with regard to these proteins are majorly restricted in isolated mutant cell strains.^76–78^ We selected 5S and TMG as the chemical tools for perturbing OGT and OGA because these compounds avoid some of the problems associated with genetic models.^79–82^ In addition, global *O*-GlcNAcylation is also regulated using siRNA against *OGT* and/or *OGA* mRNA to guarantee and compensate for the chemical regulation via mRNA suppression. Overall, these findings strongly suggest that a high level of *O*-GlcNAcylation supports progressive phenotypes of CCA cells in several aspects.

For the Click-iG strategy, 1,6-Pr_2_GalNAz can cross cell membranes, and undergo deacetylation by non-specific esterases to generate cell-active GalNAz, which can be metabolically converted to UDP-GalNAz with high efficiency in cells via the GalNAc salvage pathway.^83^ The NAD-dependent epimerase UDP-galactose-4-epimerase (GALE) converts UDP-GalNAz to UDP-GlcNAz that serves as the substrate for *O*-GlcNAcylation by the Leloir-type glycosyltransferase OGT. However, cautions are needed in glycoproteomic data interpretation, because the resulting UDP-GalNAz and UDP-GlcNAz are general nucleotide sugar donors, and can be readily incorporated into *N*-linked-, mucin-type *O*-linked-, and *O*-GlcNAcylated-glycoproteins. Indeed, the precise annotation of glycan structures is still a major challenge in intact glycopeptide analysis, so we manually annotate the *O*-GlcNAcylation site with the aid of glycoprotein subcellular location and the presence of *N*-(4-aminomethyl)-triazolylacetylglucosamine (GlcNAt).^28^ Fortunately, Click-iG enables comprehensive coverage of the protein glycosylation landscape, which provides a blueprint for interrogating the crosstalk between different glycosylation pathways. Other *O*-GlcNAc chemical reporters such as peracetylated azido- or alkynyl-modified GalNAc or GlcNAc counterparts (*i.e.*, Ac_4_GalNAz, Ac_4_GalNAlk, Ac_4_GlcNAz, Ac_3_6AlkGlcNAc) are possible candidates for Click-iG application, although non-specific S-glyco-modification should be taken into consideration.^84–86^

Keratins have long been recognized as diagnostic markers and prognostic indicators in a variety of epithelial malignancies.^34^ For instance, Keratin 19 has been identified as the CCA proliferative marker for postoperative intrahepatic cholangiocarcinoma patients and is of use in distinguishing cholangiocarcinoma from hepatocellular carcinomas.^87,88^ Our chemical glycoproteomic analysis linked Keratin 18 to its *O*-GlcNAcylation with detailed glycosite information. Nevertheless, how *O*-GlcNAcylation of K18 underscores the cellular proliferation and tumor growth has been incongruous.^89^ A previous report from the Rajiv lab indicated that *O*-GlcNAcylation decreases K18 stability and elevation of *O*-GlcNAcylation promotes ubiquitinoylation in human hepatocytes, ^49^ in contradict with our findings in this study. We verify that the removal of *O*-GlcNAc modification at Ser 30 of K18 negatively affected CCA cell proliferation and xenograft tumor growth, adding a novel mechanistic insight into the regulation of K18, and highlighting the potential to intervene in K18 *O*-GlcNAcylation as a therapeutic strategy against CCA. Doubtless, site-specific phosphorylation of K18 is well characterized for its biological roles.^90,91^ The reciprocal interplay between *O*-GlcNAcylation and phosphorylation is also a critical event in modulating protein-protein interaction, subcellular localization and protein degradation.^92–94^ Competitive site blocking between *O*-GlcNAcylation and phosphorylation (*e.g.*, Ser 49), as well as their synergism (*e.g.*, Ser 31 *O*-GlcNAcylation and Ser 34 phosphorylation), poise functional modulation in K18 solubility, filament organization, and stability.^44,95,96^ It will be interesting to investigate whether K18 *O*-GlcNAcylation has crosstalk with phosphorylation or other PTM.

The cycling of *O*-GlcNAc on nucleocytoplasmic proteins is tightly associated with glucose metabolism and nutrient flux, which is often dysregulated and reprogrammed in cancer.^50,97,98^ A recent report emphasized the down-regulation proteins in the glycolysis/gluconeogenesis pathway of large-duct type intrahepatic cholangiocarcinoma.^62^ Here we identified a previously unknown mechanism for the regulation of the TCA cycle through *O*-GlcNAc modification on K18, which interacts with TCA enzymes, namely IDH2, IDH3A, IDH3B and IDH3G. Deregulation of IDHs catalytic activities is associated with cholangiocarcinoma and other human diseases. The heterozygous gain-of-function mutation in IDHs, which catalyzes the oxidation of isocitrate to α-ketoglutarate (α-KG), generates oncometabolite 2-hydroxyglutarate (2-HG), by NADPH reduction of α-KG. 2-HG competitively inhibits the α-KG-dependent dioxygenases that catalyze the transformation from α-KG to succinate, resulting in dysregulation of epigenetic and gene expression profiles, apoptosis resistance and increased migration and invasion. Therefore, *IDHs* are considered as proto-oncogenes in cancer metabolic derangement.^97,99–101^ Given the dependence of CCA and other cancer cells on *O*-GlcNAc, targeting OGT and other components of the TCA cycle may reduce the aggressiveness of CCA cells while sensitizing them to chemotherapeutic drugs.

In summary, we find that *O*-GlcNAcylation is dysregulated in human cholangiocarcinoma, majorly attributed to the up-regulation of OGT. Global up-regulation of *O*-GlcNAcylation benefits the expression of cell cycle checkpoints such as BUB1, FOXM1 and cMyc, enhancing cell cycle progression and cell growth. To elucidate the function of *O*-GlcNAcylation in CCA progression, we used chemical glycoproteomic analysis Click-iG for systematic profiling of intact glycopeptides in CCA and HIBEpiC cell lines. *O*-GlcNAcylation of K18 at Ser 30 increases its stability, while the orchestration between glycosylated K18 and IDH proteins in the TCA cycle reveals a positive regulation of CCA progression. How knowledge of *O*-GlcNAcylation has led to the aberrant state of cell proliferation and tumorigenesis in cholangiocarcinoma will be interesting to pursue.

## Limitations of this study

Cellular metabolism of 1,6-Pr_2_GalNAz inevitably alters the endogenous level of UDP-GlcNAc and UDP-GalNAc, therefore, it is not ruled out that UDP-GlcNAz or UDP-GalNAz may interfere with the catalytic activity of glycosyltransferases in CCA cell lines. Due to the accessibility of bioorthogonal reaction and the dynamic feature of *O*-GlcNAc modification, it is still challenging to bypass the abundance bias and generate MS spectra for glycosite identification, with precise annotation. As *O*-GlcNAcylation occurs on thousands of protein substrates, targeting OGT and OGA using either chemical or biological methodologies might compromise normal biological processes. In addition, other functional consequences of K18 *O*-GlcNAcylation at other sites still remain ambiguous, because only a limited number of tools exist to study its site-specific *O*-GlcNAcylation effects.^102^ Further preclinical and clinical studies are needed to confirm the trueness of CCA-associated *O*-GlcNAcylated proteins as diagnosis/prognosis indicators. The integration of glycomics and other “omics” such as genomics or transcriptomics in CCA cell lines or tissues from patients, will provide an avenue for greater impact on developing therapeutics. Additional studies are required to address these questions in the future.

## Acknowledgments

We thank Prof. Dr. Xing Chen (Peking University) for his guidance with glycoproteomics and insightful discussions. The LC-MS/MS experiments were performed at the National Center for Protein Sciences (Beijing, China). We gratefully acknowledge support from the National Natural Science Foundation of China (2207070006 and 22107005), the Natural Science Foundation of Jiangsu Province (No. BK20202299), and the Programs for High-level Entrepreneurial and Innovative Talents Introduction of Jiangsu Province (Individual and Group Program), the Fundamental Research Funds for the Central Universities (021414380508), and the STI2030-Major Projects (2022ZD0211804).

## Author Contributions

R.X., J.L., and X.M. conceptualized and designed the experiments; R.X., J.L., and L.W. supervised the project; X.M., Y.Z., C.W., X.T., and X.Z. performed the experiments; X.M., and Y.Z. analyzed the data; L.X., and Y.Z. collected the clinical samples; B.C., and Y.H. provided materials support; R.X., J.L., and X.M. wrote this paper. All authors read and approved the final manuscript.

## Inclusion and diversity

We support inclusive, diverse, and equitable conduct of research.

## STAR ★ Methods

Detailed methods are provided in the online version of this paper and include the following:

## Key Resources Table

- **Resource Availability**
  - Lead contact
  - Materials availability
  - Data and code availability
- **Experimental Model and Study Participant Details**
  - Patient samples
  - Cell lines and cell culture
  - Animals
- **Method Details**
  - Immunohistochemistry (IHC) analysis
  - Protein extraction and Western blot analysis
  - RNA extraction and quantitative real-time PCR (qRT-PCR)
  - Cell counting kit-8 (CCK-8) assay
  - Cell apoptosis assay
  - Cell cycle analysis
  - Colony formation assay
  - Metabolic oligosaccharide engineering (MOE) of living cells
  - Glycopeptides enrichment
  - LC-MS/MS analysis and data processing
  - Chemoenzymatic labeling of *O*-GlcNAcylated proteins
  - Immunoprecipitation (IP) assay
  - *O*-GlcNAcylation stoichiometry on K18
  - Plasmids construction
  - Lentivirus infection and stable cell line establishment
  - Determination of K18 half-life
  - Ubiquitination assay
  - Tumorigenesis in nude mice
  - Analysis of *O*-GlcNAcylated K18 in tissues
  - Interacting proteins analysis
  - Analysis of metabolites by LC–MS/MS
- **Quantification and Statistical Analysis**

## STAR★Methods

### Key Resources Table

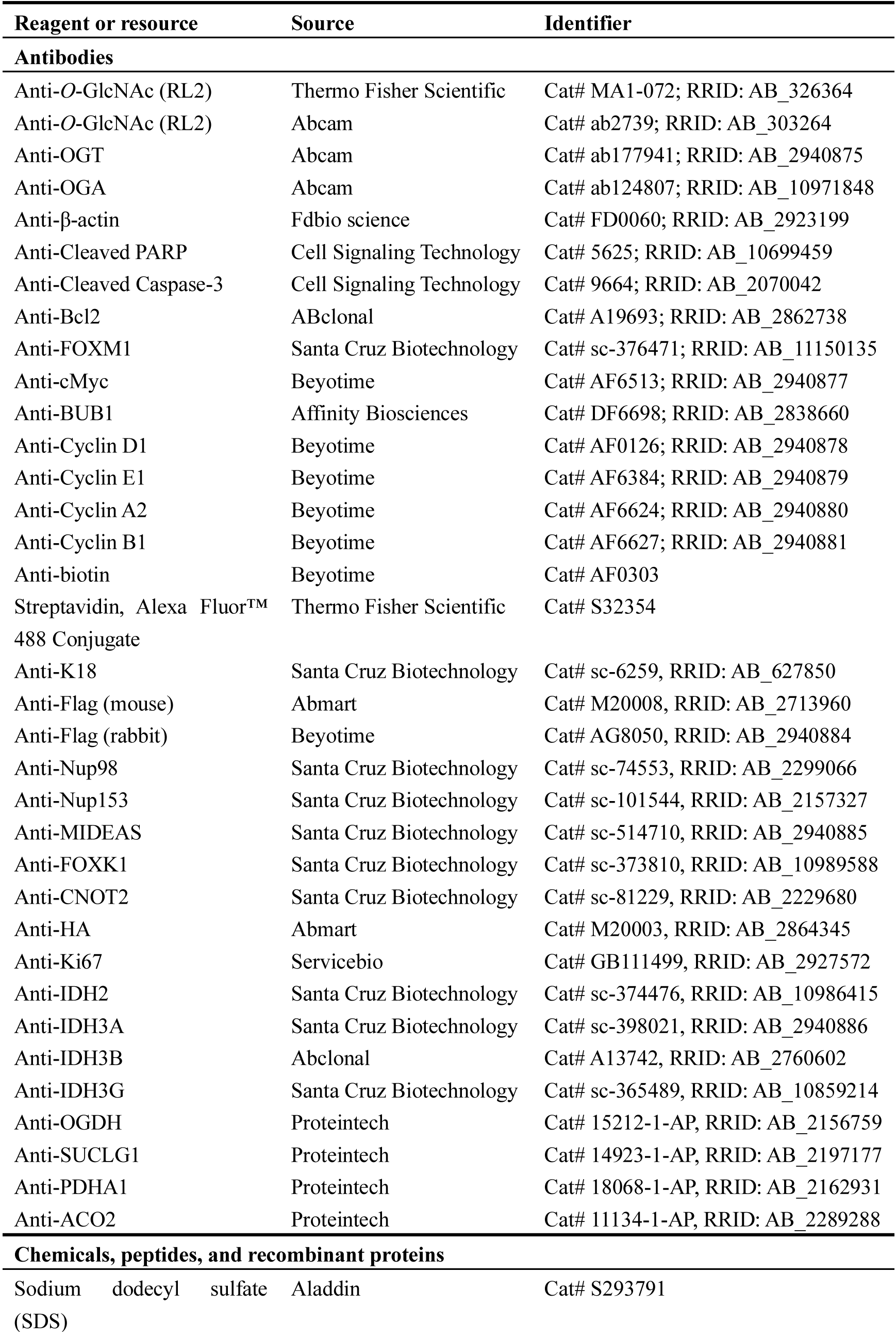
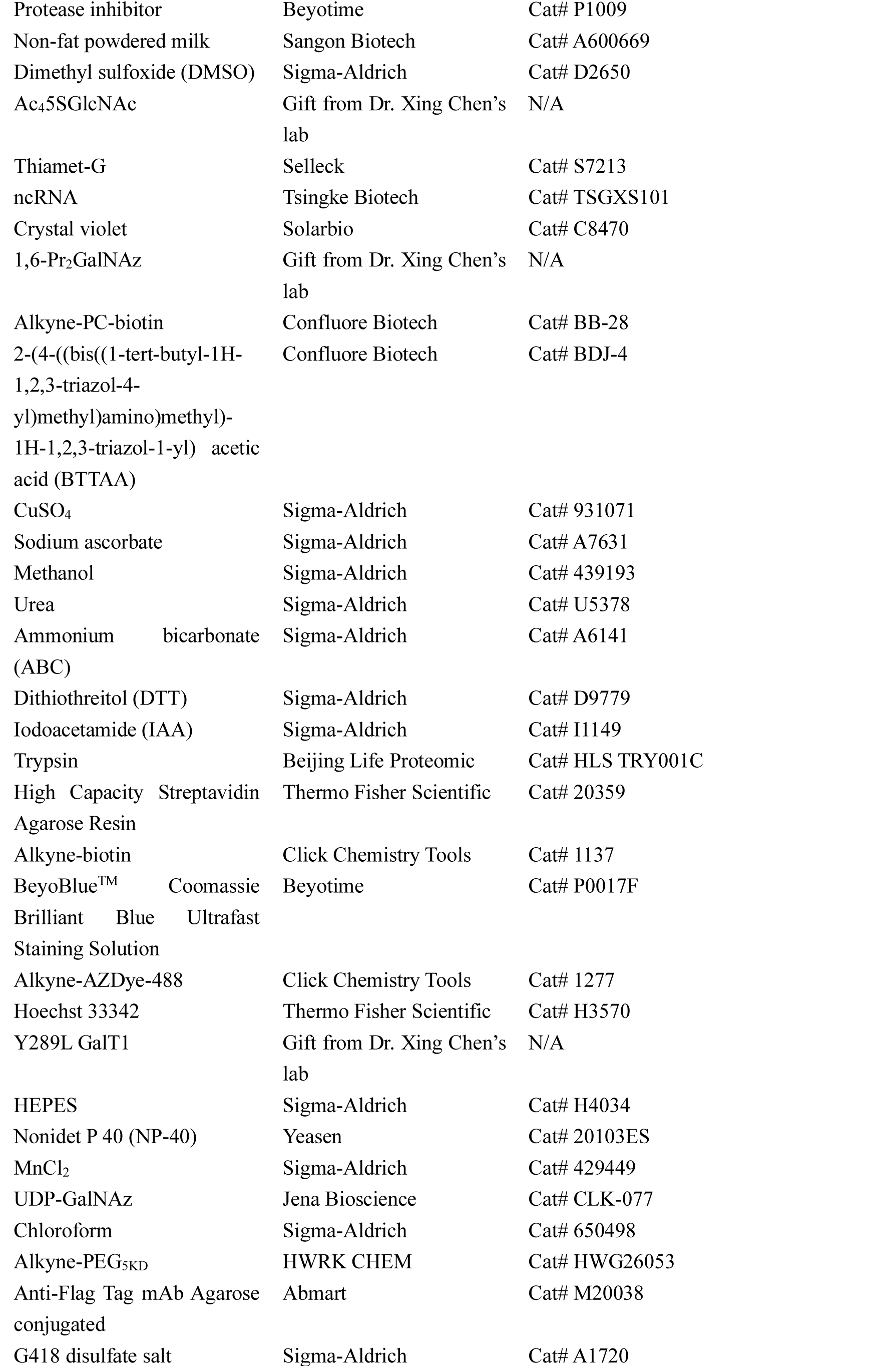
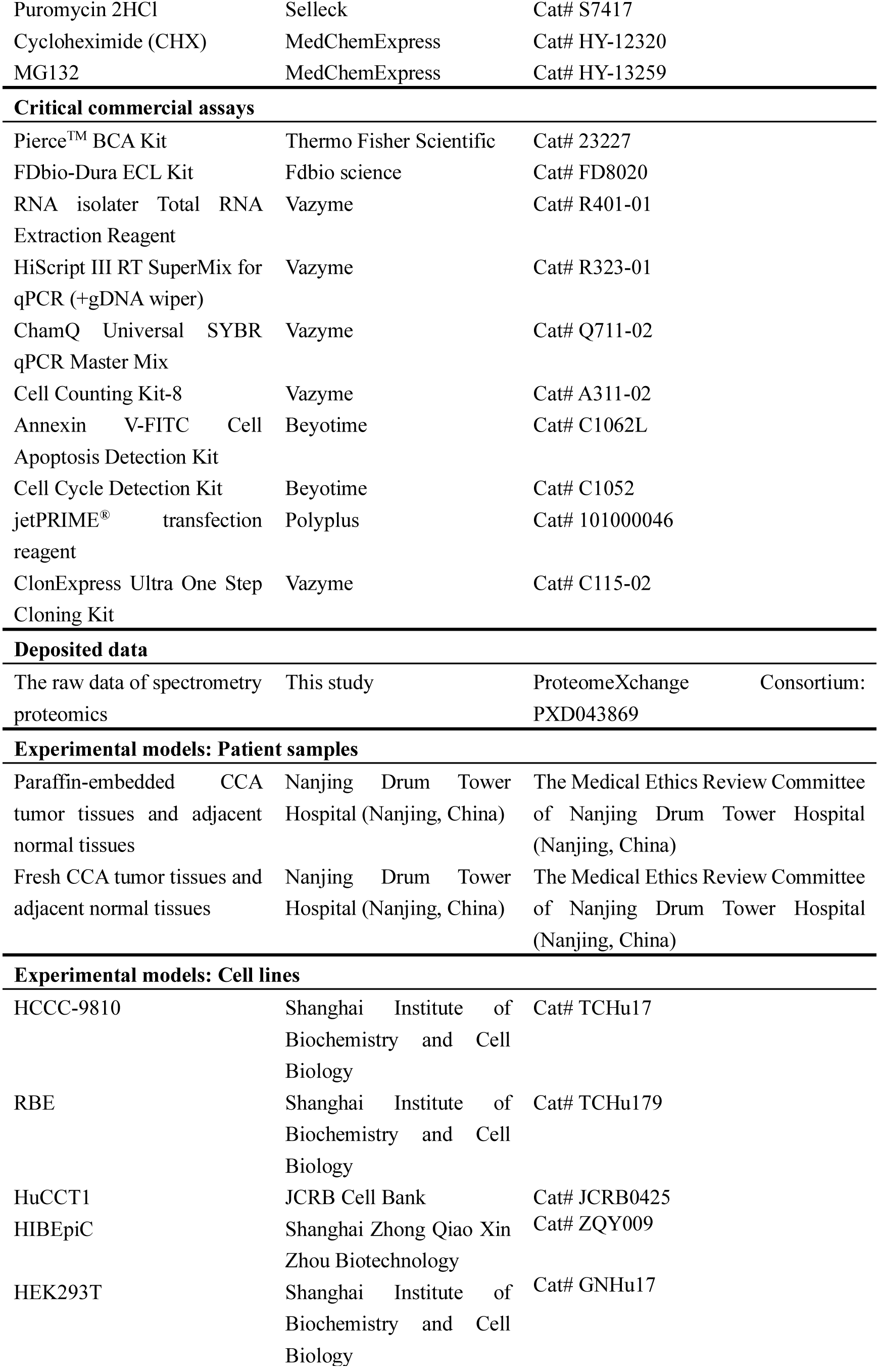
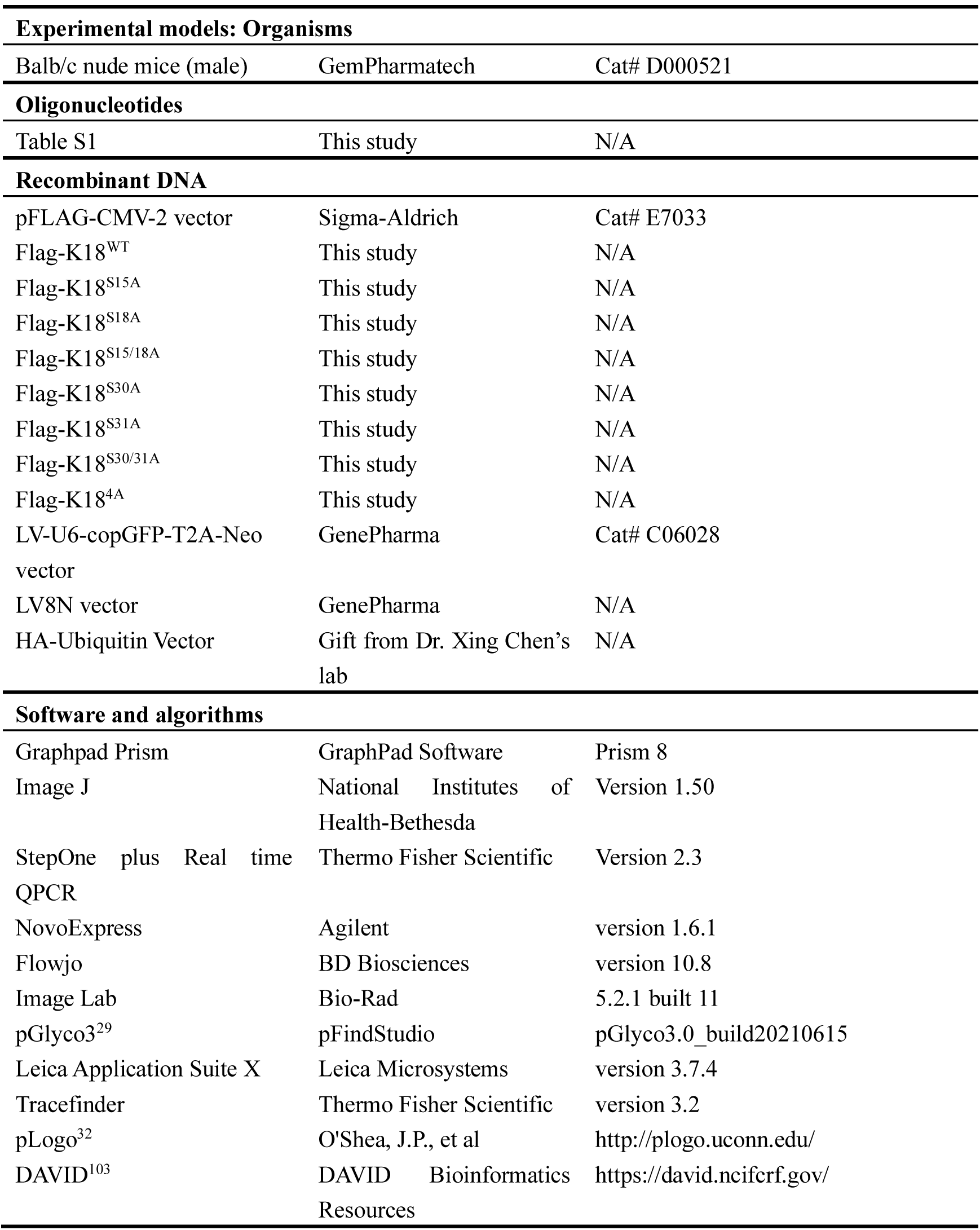

### Resource Availability

#### Lead contact

Further information and requests for resources and reagents should be directed to and will be fulfilled by Dr. Ran Xie.

#### Materials availability

Materials generated in this study are accessible upon request.

#### Data and code availability

The mass spectrometry data have been deposited to ProteomeXchange Consortium (http://proteomecentral.proteomexchange.org), and are publicly available as of the date of publication. Accession numbers are listed in the key resources table.

This paper does not report the original code.

Any additional information required to reanalyze the data reported in this paper is available from the lead contact upon request.

### Experimental Model and Study Participant Details

#### Patient samples

Human cholangiocarcinoma (CCA) tumor tissues and adjacent normal tissues were collected from patients undergoing surgery at the Drum Tower Hospital Affiliated to the Medical School of Nanjing University (Nanjing, China). Written consent was obtained from all patients, and all experiments in this study were conducted in accordance with official guidelines (Declaration of Helsinki), approved by the Medical Ethics Review Committee of Nanjing Drum Tower Hospital (Nanjing, China).

#### Cell lines and cell culture

Human CCA cells HCCC-9810 and RBE, and HEK293T cells were purchased from the Institute of Biochemistry and Cell Biology, Shanghai Institutes for Biological Sciences, Chinese Academy of Sciences, Shanghai, China. Human CCA cells HuCCT1 were purchased from the Japanese Collection of Research Bioresources Cell Bank (JCRB, Osaka, Japan). Human Intrahepatic Biliary Epithelial Cells (HIBEpiC) were purchased from Shanghai Zhong Qiao Xin Zhou Biotechnology (Shanghai, China). HCCC-9810, RBE, and HuCCT1 cells were cultured in RPMI 1640 medium (Gibco, CA, USA). HEK293T cells were cultured in DMEM medium (Gibco, CA, USA). HIBEpiC cells were cultured in Epithelial Cell Medium supplementing with 1% Epithelial Cell Growth Supplement (EpiCGS), which was purchased from Shanghai Zhong Qiao Xin Zhou Biotechnology (Shanghai, China). All cell lines were cultured with 10% fetal bovine serum (FBS, Gibco, USA), and 100 U/mL of penicillin and streptomycin (Gibco, USA) in a cell incubator at 37 ℃ with 5% CO_2_. All cell lines were tested negative for mycoplasma contamination. Cells were transfected with small interfering RNA (siRNA) or plasmid in this study using jetPRIME^®^ transfection reagent (101000046, Polyplus) when they reached approximately 80% confluence.

#### Animals

5-week-old BALB/c nude mice were purchased from GemPharmatech (Nanjing, China) and housed under specific pathogen-free (SPF) conditions. All animal experimental procedures were approved by the Animal Care and Use Committee of Nanjing University.

### Method Details

#### Immunohistochemistry (IHC) analysis

The formalin-fixed, paraffin-embedded samples were sliced, dewaxed, and rehydrated, followed by incubation with anti-*O*-GlcNAc (RL2, 1:200, MA1-072, Thermo Fisher Scientific), anti-OGT (1:200, ab177941, Abcam), anti-OGA (1:200, ab124807, Abcam) or anti-Ki67 (1:200, GB111499, Servicebio) antibody, and secondary antibody. After that, samples were incubated with diaminobenzidine (DAB) (Dako, USA) and restained with hematoxylin (Sigma-Aldrich). The analysis of the IHC score (RL2 (anti-*O*-GlcNAc), OGT and OGA staining) of CCA tumor tissues and adjacent normal bile duct (BD) was performed as previously described.^62^ Level of staining: 0-negative, 1-weakly positive, 2-positive, 3-strongly positive. For Ki-67 staining, the positive area of xenograft tumors was determined through IHC analysis.

#### Protein extraction and Western blot analysis

The cells were lysed using 4% sodium dodecyl sulfate (SDS, wt/vol) supplementing with protease inhibitor (Beyotime, Shanghai, China) under sonication. After centrifugation, the supernatant was quantified for protein concentration using a bicinchoninic acid (BCA) kit (Pierce, USA).

For Western blot analysis, cell lysates supplemented with 5× loading buffer were boiled at 99 ℃ for 5 min. Equal amounts of proteins were separated by SDS-PAGE gel and blotted onto a polyvinylidene fluoride (PVDF) membrane. After blocking with 5% non-fat powdered milk (wt/vol), the membrane was incubated with specific primary antibodies for anti-*O*-GlcNAc (RL2, 1:2000, ab2739, Abcam), anti-OGT (1:2000, ab177941, Abcam), anti-OGA (1:2000, ab124807, Abcam), anti-β-actin (1:5000, FD0060, Fdbio), anti-Cleaved Caspase-3 (1:1000, #9664, Cell Signaling Technology), anti-Bcl2 (1:1000, A19693, ABclonal), anti-FOXM1 (1:1000, sc-376471, Santa Cruz), anti-cMyc (1:1000, AF6513, Beyotime), anti-BUB1 (1:1000, DF6698, Affinity), anti-Cyclin D1 (1:1000, AF0126, Beyotime), anti-Cyclin E1 (1:1000, AF6384, Beyotime), anti-Cyclin A2 (1:1000, AF6624, Beyotime), anti-Cyclin B1 (1:1000, AF6627, Beyotime), anti-K18 (1:1000, sc-6259, Santa Cruz), anti-FLAG (1:2000, M20008, Abmart), anti-Nup98 (1:1000, sc-74553, Santa Cruz), anti-Nup153 (1:1000, sc-101544, Santa Cruz), anti-MIDEAS (1:1000, sc-514710, Santa Cruz), anti-FOXK1 (1:1000, sc-373810, Santa Cruz), anti-CNOT2 (1:1000, sc-81229, Santa Cruz), anti-HA (1:1000, M20003, Abmart), anti-IDH2 (1:1000, sc-374476, Santa Cruz), anti-IDH3A (1:1000, sc-398021, Santa Cruz), anti-IDH3B (1:1000, A13742, ABclonal), anti-IDH3G (1:1000, sc-365489, Santa Cruz), anti-OGDH (1:1000, 15212-1-AP, Proteintech), anti-SUCLG1 (1:1000, 14923-1-AP, Proteintech), anti-PDHA1 (1:1000, 18068-1-AP, Proteintech) and anti-ACO2 (1:1000, 11134-1-AP, Proteintech), followed by secondary horseradish peroxidase (HRP)-conjugated antibodies. After washes with Tris-buffered saline with Tween 20 (TBST), blots were reacted with enhanced chemiluminescence (ECL) reagent (Fdbio science) and protein bands were detected by a chemiluminescence system (Tanon-5200, Shanghai, China).

#### RNA extraction and quantitative real-time PCR (qRT-PCR)

Total RNA was isolated from cultured cells by RNA isolater total RNA extraction reagent (Vazyme, Nanjing, China) according to the manufacturer’s instructions. To quantify mRNAs, total RNA was converted to cDNA using the reverse transcription kit (Vazyme), followed by PCR using ChamQ universal SYBR qPCR master mix reagent (Vazyme) and gene-specific primers. All of the reactions were run in triplicate. The expression levels of mRNAs were normalized to *ACTB* mRNA using the 2^-ΔΔCт^ method. The primers used for qRT-PCR were provided in the key resources table (**Table S1**).

#### Cell counting kit-8 (CCK-8) assay

The cell viability of HuCCT1, RBE and HCCC-9810 cells was determined using the CCK-8 assays (Vazyme) following the manufacturer’s instructions. Briefly, HuCCT1, RBE and HCCC-9810 cells were seeded into 96-well plates at a density of 1×10^4^ cells/well, followed by exposure to a gradient concentration of Ac_4_5SGlcNAc (5S) and Thiamet-G (TMG) for 48 h. Then, 100 μL/well RPMI 1640 medium containing 10% CCK-8 was replaced into the test well and incubated at 37 ℃ for 1 h. Absorbance was then measured at a wavelength of 450 nm.

#### Cell apoptosis assay

HuCCT1, RBE and HCCC-9810 cells were seeded into 6-well plates at a density of 4×10^5^ cells/well, followed by exposure to a gradient concentration of 5S and TMG. After incubation for 48 h, the cells were harvested for apoptosis analysis. Briefly, the cells were washed twice with cold PBS and resuspended in 1×binding buffer with 1×10^6^ cells/mL, following FITC-Annexin V (FITC fluorescence) and PI (PE fluorescence) were added. The cells were incubated at room temperature for 15 mins in the dark and were analyzed by flow cytometry (Agilent) within 1 h after staining.

#### Cell cycle analysis

Cell cycle distribution of HuCCT1, RBE and HCCC-9810 cells followed by exposure to a gradient concentration of 5S and TMG for 48 h was detected by FACS analysis. Briefly, cells were trypsinized into single-cell suspension, rinsed with ice-cold PBS and fixed in ice-cold 70% ethanol overnight. Then the cells were stained in PI added with RNase A (100 μg/ml) at 37 ℃ for 30 mins, and determined by flow cytometry (Agilent).

#### Colony formation assay

HuCCT1 and RBE cells transfected with ncRNA, siOGT-1, siOGT-2, siOGT-3, siOGA-1, siOGA-2 or siOGA-3 were seeded into 6-well plates at a density of 300 cells/well, and cultured in RPMI 1640 medium supplemented with 10% FBS for 14 days, during which the medium was replaced every 3 days. Colonies were then fixed with methanol for 10 mins and stained with 4% crystal violet in PBS for 15 mins. Colony formation was shown by the number of stained colonies.

#### Metabolic oligosaccharide engineering (MOE) of living cells

HuCCT1, HIBEpiC, HCCC-9810 and RBE cells seeded at 10-cm dishes were treated with unnatural sugar (1,6-Pr_2_GalNAz) at varied concentrations for 48 h or with 200 μM 1,6-Pr_2_GalNAz for up to 72 h when they reached approximately 30% confluence. The cells were harvested by trypsin digestion and washed twice with PBS, and were lysed as described in the ‘Protein extraction and Western blot analysis’ section. All lysates were incubated with 500 μM premixed CuSO_4_/BTTAA (molar ratio 1:2), 100 μM alkyne-biotin and 2.5 mM fresh sodium ascorbate for 2 h at room temperature, followed adding with 5×loading buffer were boiled at 99 ℃ for 5 min. Equal amounts of proteins were detected with anti-biotin (1:2000, A0303, Beyotime) by Western blot analysis. For confocal fluorescence microscopy imaging, the cells were seeded into 8-chamber at 1×10^4^ cells/well and treated with 200 μM 1,6-Pr_2_GalNAz for 48 h. The cells were washed twice with PBS, and then fixed with 4% paraformaldehyde (wt/vol), and permeabilized with 0.5% Triton-X 100 (vol/vol). Then, the cells were incubated with 50 μM premixed CuSO_4_/BTTAA (molar ratio 1:6), 50 μM alkyne-AZDye-488 and 2.5 mM fresh sodium ascorbate for 10 mins at room temperature. For nucleus staining, cells were incubated with 5 μg/mL Hoechst 33342 at room temperature for 20 min. The cells were washed three times after each step. Finally, the cells were imaged by Leica TCS SP5 laser scanning confocal system equipped with a ×63 oil immersion objective lens. For FACS analysis, the cells were seeded at 6-well plates and treated with 200 μM 1,6-Pr_2_GalNAz for 48 h when they reached approximately 30% confluence. The cells were trypsinized into single-cell suspension, rinsed with ice-cold PBS and fixed in ice-cold 70% ethanol overnight. Then, the cells were incubated with 50 μM premixed CuSO_4_/BTTAA (molar ratio 1:6), 50 μM alkyne-biotin and 2.5 mM fresh sodium ascorbate for 10 mins on ice, following incubated with streptavidin-488 for 30 mins on ice, and determined by flow cytometry (BD Biosciences).

#### Glycopeptides enrichment

The lysates extracted from HuCCT1 and HIBEpiC cells were incubated with 500 μM premixed CuSO_4_/BTTAA (molar ratio 1:2), 100 μM alkyne-PC-biotin and 2.5 mM fresh sodium ascorbate for 3 h at room temperature. The mixture was added to 8 volumes of ice-cold methanol for precipitation overnight at -80 ℃ and washed three times with ice-cold methanol. Then, the protein pellet was reconstituted by sonication using 4 M urea in 50 mM ammonium bicarbonate (ABC), incubated with 10 mM dithiothreitol (DTT) at 37 ℃ for 1 h, followed by incubation with 20 mM iodoacetamide (IAA) at room temperature for 30 mins in the dark. The solution was diluted to 1 M urea in 50 mM ABC supplemented with mass spectra grade trypsin (enzyme: substrate ratio at 1: 50) and reacted at 37 ℃ for 16 h. After that, the streptavidin agarose beads (150 µL per 40 mg, Thermo Fisher Scientific, 20359) were added to the solution above and gently rotated for 3 h at room temperature. The beads were washed five times with PBS and Milli-Q water successively and re-suspended with 200 µL 0.1% formic acid (vol/vol), followed by irradiating three times under 365 nm UV light for 5 min using a UV cross-linker (CL-1000; UVP). The supernatant was collected, evaporated in a vacuum centrifuge, and subjected to liquid chromatography-tandem mass spectrometry (LC-MS/MS) analysis.

#### LC-MS/MS analysis and data processing

LC-MS/MS analysis of the glycopeptides enriched above was performed as previously described.^28^ In brief, all samples were resuspended with 0.1% FA, and subjected to analysis using a Dionex Ultimate 3000 RPLC nano system, which was connected with an Orbitrap Fusion Lumos Tribrid Mass Spectrometer with an EASY-Spray ionization source (Thermo Fisher Scientific). Survey scans of precursor were collected in Orbitrap from 350-2000 Th, under the resolution of 120,000 at 200 m/z. Monoisotopic precursors selection was enabled, and the multi-charged precursors with z = 2-8 were selected for data-dependent MS/MS scans with a cycle time of 3 s, dynamic exclusion set to 15 s, and window set to ±10 ppm. The initial data-dependent MS/MS scans were acquired using HCD with a first mass of 100 Th, a normalized collision energy (NCE) of 30 ±10, and a resolution of 30,000 at 200 m/z, according to the sceHCD-pd-EThcD method. Following ETD fragmentation triggered by glycan oxonium fragments, the glycan oxonium ions (*i.e*, m/z 168.0654, 186.0760, 204.0865, 300.1302, 366.1395, 388.1469, etc.) were detected in the sceHCD spectrum with a mass accuracy within 10 ppm, meanwhile, additional precursor isolation and EThcD acquisition were performed with supplemental activation of 35.

The raw data processing was performed using pGlyco3 (https://github.com/pFindStudio/pGlyco3/releases/tag/pGlyco3.0.rc3_build2021 0124) under the “HCD + EThcD” mode as previously described.^28,29^ Briefly, MS/MS spectrum were searched against the SwissPort Mus musculus proteome database downloaded from Uniprot (https://www.uniprot.org) on 4th November 2016. The *N*-glycans and *O*-glycans were searched against pGlyco-*N*-glycan mode and pGlyco-*O*-glycan mode in the pGlyco3 software with modified glycan databases, respectively. The parameter precursor tolerance was set to ± 10 ppm and fragment tolerance ± 20 ppm. The false discovery rate (FDR) was set to less than 1%. Notably, the *O*-GlcNAc sites were assigned manually based on the subcellular localization. The proteins localized in the cytoplasmic side (including the nucleus, cytoplasmic, mitochondrial and cytoplasmic part of transmembrane proteins) were selected as *O*-GlcNAc proteins. For quantification, the *O*-GlcNAc sites co-identified in three independent replicates of HuCCT1 and HIBEpiC cells were defined as quantified *O*-GlcNAc sites. The *O*-GlcNAc sites with P-value < 0.05 and a fold change > 1.50 or < 0.67 were considered as up-regulated or down-regulated *O*-GlcNAc sites, respectively.

#### Chemoenzymatic labeling of *O*-GlcNAcylated proteins

The cells were lysed as described in the ‘Protein extraction and Western blot analysis’ section, and the lysates were added to 8 volumes of ice-cold methanol overnight at -80 ℃ and washed three times with ice-cold methanol. The proteins were reconstituted with 1% SDS (wt/vol) in 20 mM HEPES buffer (pH 7.9) using sonication, and incubated with 125 mM NaCl, 5% Nonidet P 40 (NP-40, vol/vol), 50 mM HEPES (pH 7.9), 100 mM MnCl_2_, 500 μM UDP-*N*-azidoacetylglucosamine (UDP-GalNAz), and a β-1,4-galactosyltransferase mutant (Y289L GalT1) (enzyme: substrate ratio at 1: 40) at 4 ℃ for 20 h. Methanol, chloroform and Milli-Q water are successively added to the solution above (solution: methanol: chloroform: Milli-Q water ratio at 1: 3: 0.75: 2) to obtain protein pellet, following washed three times with ice-cold methanol. The proteins were resuspended as described above.

#### Immunoprecipitation (IP) assay

For biotin immunoprecipitation, the cells were labeled and lysed as mentioned above. The biotinylated lysates were treated with streptavidin agarose beads (10 µL/mg), and gently rotated for 3 h at room temperature. For FLAG immunoprecipitation, the cells transfected with indicated plasmids were labeled and lysed as mentioned above. Then, the biotinylated lysates were treated with anti-FLAG beads (10 µL/mg, M20038, Abmart) at 4 ℃ for 12 h with gentle rotation. The beads were washed five times with PBS, following added with 5×loading buffer and boiled at 99 ℃ for 5 min. The final immunoprecipitated proteins were analyzed by western blot.

#### *O*-GlcNAcylation stoichiometry on K18

The cells treated with metabolic oligosaccharide engineering or chemoenzymatic labeling were incubated with 500 μM premixed CuSO_4_/BTTAA (molar ratio 1:2), 100 μM alkynylated polyethylene glycol 5000 (alkyne-PEG_5KD_, HWRK CHEM, Beijing, China) and 2.5 mM fresh sodium ascorbate at 37 ℃ for 16 h, followed adding with 5×loading buffer were boiled at 99 ℃ for 5 min. Equal amounts of proteins were detected with anti-K18 (Santa Cruz) by Western blot analysis.

#### Plasmids construction

The FLAG-tagged K18 constructs were generated by in-frame subcloning the human *KRT18* cDNA into the pFLAG-CMV-2 vector (Sigma-Aldrich). The K18 mutants (S15A, S18A, S15/18A, S30A, S31A, S30/31A and 4A) were generated using the ClonExpress Ultra One Step Cloning Kit (C115-02, Vazyme) according to the manufacturer’s protocol. All constructs were confirmed by DNA sequencing (Sangon, Shanghai, China). All primers used in plasmids construction can be found in the key resources table (**Table S1**).

#### Lentivirus infection and stable cell line establishment

The lentiviruses with random small hairpin RNA (shNC, used as a negative control) and small hairpin RNA K18 knockdown (shK18-1∼3) were generated by GenePharma (Shanghai, China) using the LV-U6-copGFP-T2A-Neo vector (GenePharma) according to the manufacturer’s instructions. The HuCCT1 cells were infected with these lentiviruses and selected for stable cell lines with 400 μg/mL G418 for two weeks. We screened the lentivirus with the highest efficiency of knockdown of endogenous K18 (shK18) by qRT-PCR and western blot analysis. To generate K18 reconstituted stable cell lines, we used the co-expressing exogenous FLAG-K18 WT or FLAG-K18 S30A lentivirus which was constructed with LV8N vector (Mock, GenePharma) to infect the shK18 HuCCT1 cells and selected for stable cell lines with 2 μg/mL puromycin for two weeks. The successful construction of stable cell lines shNC + Mock, shK18, shK18 + WT and shK18 + S30A were verified by qRT-PCR and western blot analysis. For CCK-8 analysis, the cells were seeded into 96-well plates at a density of 5×10^3^ cells/well, and the viability of cells was determined using the CCK-8 assays (Vazyme) following the manufacturer’s instructions. Cell cycle distribution of the cells above was by FACS analysis as described in the ‘Cell cycle analysis’ section. For colony formation, the cells were seeded into 6-well plates at a density of 200 cells/well, and detected as described in the ‘Colony formation assay’ section.

#### Determination of K18 half-life

The degradation of K18 in HuCCT1, RBE and HCCC-9810 cells incubated with DMSO, 200 μM 5S or 1 μM TMG for 48 h, followed by treatment with 10 μg/mL cycloheximide (CHX, for 0, 2,4 or 8 h) was determined by Western blot analysis. The stable cell lines (shK18 + WT and shK18 + S30A) were treated with 10 μg/mL CHX for 0 or 8 h. The FLAG-K18 protein levels were analyzed by western blot.

#### Ubiquitination assay

The stable cell lines (shK18 + WT and shK18 + S30A) were transfected with HA-ubiquitin and treated with 0 or 5 μM MG-132 for 20 h. Then, the cells were lysed and captured with anti-FLAG beads. Anti-HA blot demonstrated the ubiquitination of immunoprecipitated FLAG-K18.

#### Tumorigenesis in nude mice

All animal experimental procedures were approved by the Animal Care and Use Committee of Nanjing University. Specifically, 5-week-old BALB/c nude mice as mentioned above were randomly divided into four groups (n = 6 per group) and injected subcutaneously with 5×10^6^ HuCCT1 stable cells with shNC + Mock, shK18, shK18 + WT and shK18 + S30A K18. shNC + Mock was the control group. Tumors generated by xenograft HuCCT1 stable cells were measured every six days from day 10. After 34 days, tumors were dissected, photographed, weighed, and subjected to immunohistochemistry analysis of the Ki67 protein.

#### Analysis of *O*-GlcNAcylated K18 in tissues

The tissues of xenograft tumors generated from HuCCT1 cells (shK18 + WT and shK18 + S30A) and patient samples were ground to single cells and lysed as described in the ‘Protein extraction and Western blot analysis’ section. Then, the lysates were biotinylated by chemoenzymatic labeling, and captured with streptavidin beads. The *O*-GlcNAcylation of K18 was determined by Western blot analysis.

#### Interacting proteins analysis

The lysates from shK18 cells transfected with FLAG-K18^WT^ or FLAG-K18^S30A^ were captured with anti-FLAG beads and analyzed by LC-MS/MS. For protein quantification, the fold change from three biological replicates of the interacting proteins with K18 in FLAG-K18^WT^ cells was calculated relative to FLAG-K18^S30A^ cells. Proteins with P-value < 0.05 and a fold change > 1.50 or < 0.67 were considered as up-regulated or down-regulated interacting proteins, respectively. Kyoto Encyclopedia of Genes and Genomes (KEGG) enrichment analysis for the upregulated interacting proteins was performed using DAVID Bioinformatics Resources (https://david.ncifcrf.gov/).

#### Analysis of metabolites by LC–MS/MS

The extraction and analysis of cell metabolic products were carried out according to the previous procedures.^104^ Briefly, the medium of cells seeded at the 10-cm dish was absorbed completely by a pump. The cells were rinsed with pre-cooled PBS for three times, and the dish was immediately placed on dry ice with 80% pre-cooled methanol (vol/vol), and incubated at -80 °C for 1 h. Subsequently, the cells were scraped off on dry ice using a cell scraper, and the lysates were centrifuged at 14,000 g for 20 mins at 4 °C to obtain the supernatant-containing metabolites. The pellets were lysed and quantified as described in the ‘Protein extraction and Western blot analysis’ section, and the supernatant containing metabolites was dried into powder using SpeedVac and lyophilize. The samples were resuspended using 100 μL 50% methanol (vol/vol). After centrifugation, 1μL supernatant was injected into the Q Exactive mass spectrometer (Thermo, CA) for detection. For each metabolite, the standard compound was detected to ensure proper chromatographic elution time and generate a standard curve. LC-MS/MS analysis was performed as previously described.^105^ Data were acquired and processed using Tracefinder software. The quantity of the metabolite fraction was normalized to the corresponding protein.

### Quantification and Statistical Analysis

Data from three independent experiments are shown as the mean ± standard deviation (SD). All software used in this study can be found in the key resources table. Student’s t-tests (two-tailed) were used to compare two data sets, and a P-value of < 0.05 was considered statistically significant (*P < 0.05, **P < 0.01, ***P < 0.001, ****P < 0.0001, ns, not significant). Otherwise, the P-value of the Kaplan-Meier survival curve (**Figures S2C and S2D**) was analyzed by the log-rank (Mantel-Cox) test, which is a hypothesis test to compare the survival distributions of two samples. The adjusted P-value (adjusted by a conservative Bonferroni correction) of motif analysis in *O*-GlcNAc sites, *N*-glycosites and mucin-type *O*-glycosites was analyzed using the pLogo (https://plogo.uconn.edu/) (**Figures S9B and S10D**). Gene ontology terms (**Figures S9C**) and the KEGG enrichment analysis (**Figures 6B**) were performed using DAVID Bioinformatics Resources (https://david.ncifcrf.gov/) and the enrichment P-values were adjusted by a modified Fisher’s exact test.

