## Supplemental Information for "*O*-GlcNAcylation of Keratin 18 coordinates TCA cycle to promote cholangiocarcinoma progression"

**Figure S1, related to Figure 1C**

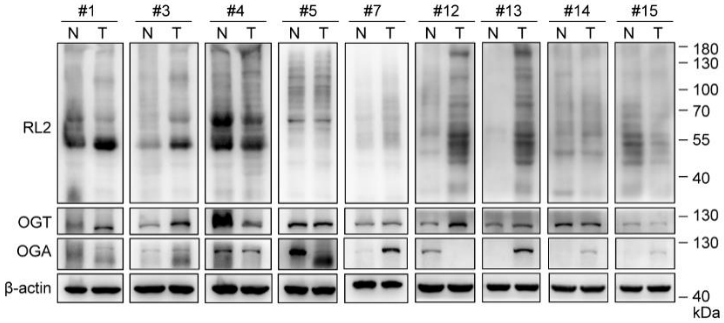


**Figure S1. *O*-GlcNAcylation is dysregulated in human CCA, related to Figure 1C.** Representative images of *O*-GlcNAcylated proteins, OGT and OGA levels from nine CCA tumor tissues (T) and adjacent normal tissues (N) by Western blot analysis. Equal loading was confirmed using β-actin. CCA, cholangiocarcinoma; *O*-GlcNAc, β-*O*-linked *N*-acetylglucosamine; RL2, anti-*O*-GlcNAc; *O*-GlcNAcylation, *O*-linked β-*N*-acetylglucosaminylation; OGT, *O*-GlcNAc transferase; OGA, *O*-GlcNAcase.

**Figure S2, related to Figure 1**

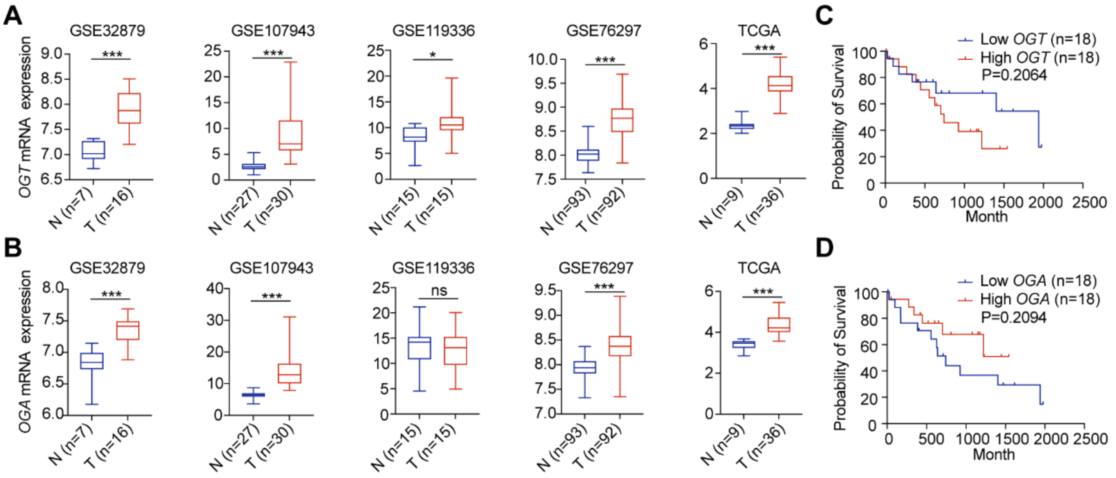


**Figure S2. *OGT* and *OGA* mRNA expression is dysregulated in CCA and correlates with OS, related to Figure 1.**

**(A)** The *OGT* and **(B)** *OGA* mRNA expression in CCA tumor tissues and adjacent normal tissues base on GSE32879, GSE107943, GSE119336, GSE76297 and TCGA datasets.

**(C)** Kaplan-Meier survival curve in the high and low *OGT* or **(D)** *OGA* mRNA expression groups. OS, overall survival.

Data (**Figures S2A and S2B**) were shown as the mean ± standard deviation (SD); statistical significance was determined by Student's t-tests (two-tailed, *P < 0.05, ***P < 0.001, ns, not significant). The P-value of the Kaplan-Meier survival curve (**Figures S2C and S2D**) was analyzed by log-rank (Mantel-Cox) test.

### Figure S3, related to Figure 2

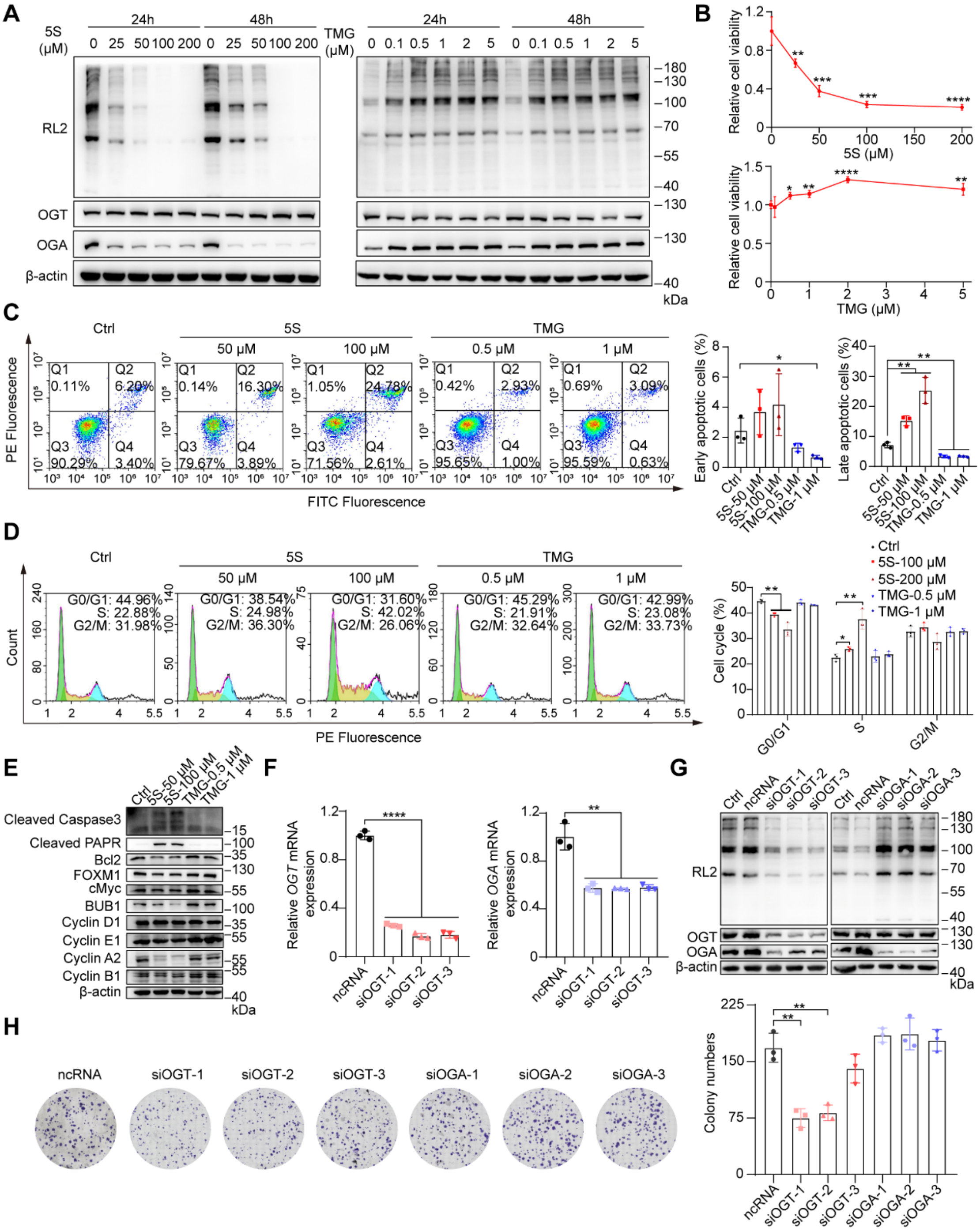

**Figure S3. *O*-GlcNAcylation promotes cell proliferation in RBE cells, related to Figure 2.**

**(A)** Time and dose-dependent analysis of the *O*-GlcNAcylated protein, OGT and OGA levels in 5S ($Ac_4$5SGlcNAc, an inhibitor of OGT)- or TMG (Thiamet-G, an inhibitor of OGA)-treated RBE cells by Western blot. Equal loading was confirmed using β-actin.

**(B)** Cytotoxicity assay of 5S- or TMG-treated RBE cells. The cells were treated with 5S or TMG at various concentrations for 48 h.

**(C)** Cell apoptosis assay of 5S- or TMG-treated RBE cells. The bivariate density plot in flow cytometry indicated the cell population of early apoptotic cells ($FITC^+/PE^-$) and late apoptotic cells ($FITC^+/PE^+$). Quantitative analysis was shown in the right panel.

**(D)** Cell cycle distribution assay of 5S- or TMG-treated RBE cells. Histogram plot in flow cytometry indicated the percentage of cell populations in the G0/G1, S, or G2/M phase. Quantitative analysis was shown in the right panel.

**(E)** Cell cycle and apoptosis marker analysis of 5S- or TMG-treated RBE cells by Western blot. Protein level of cleaved Caspase3, cleaved PARP (apoptotic markers); Bcl2 (anti-apoptotic marker); cMyc, Cyclin D1, FOXM1, Cyclin E1(G1/S transition markers); BUB1, Cyclin A2, Cyclin B1(G2/M transition markers) were analyzed. Equal loading was confirmed using β-actin.

**(F)** The *OGT* or *OGA* mRNA expression analysis of small interfering RNA (siRNA)-treated RBE cells by qRT-PCR. Random RNA was used as the negative control (ncRNA). Three independent siRNAs targeting *OGT* (siOGT1-3) or *OGA* (siOGA1-3) were shown.

**(G)** Silencing efficacy of siOGT or siOGA in RBE cells. the *O*-GlcNAcylated protein, OGT and OGA levels were analyzed by Western blot. Equal loading was confirmed using β-actin.

**(H)** Clonogenic assay of RBE cells transfected with ncRNA, siOGTs or siOGAs. Colony numbers were quantified in the right panel.

Data were shown as the mean ± SD; statistical significance was determined by Student's t-tests (two-tailed, *P < 0.05, **P < 0.01, ***P < 0.001, ****P < 0.0001).

**Figure S4, related to Figure 2**

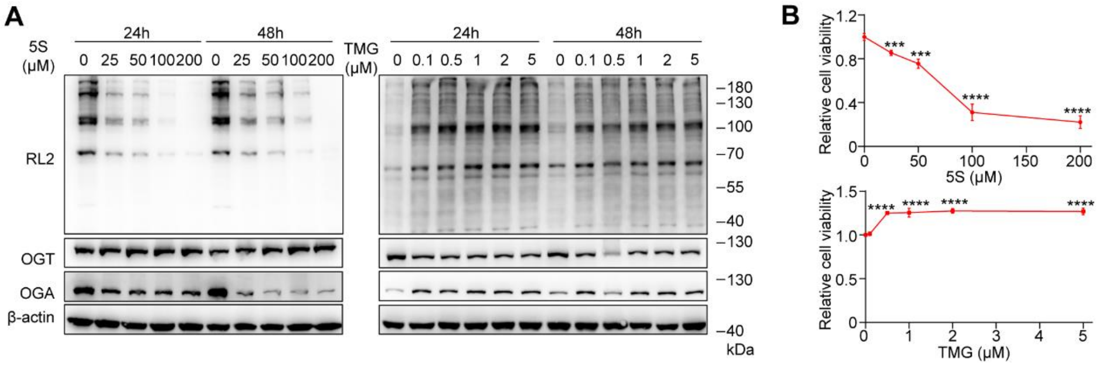


**Figure S4. *O*-GlcNAcylation promotes cell proliferation in HCCC-9810 cells, related to Figure 2.**

**(A)** Time and dose-dependent analysis of the *O*-GlcNAcylated protein, OGT and OGA levels in 5S- or TMG-treated HCCC-9810 cells by Western blot. Equal loading was confirmed using β-actin.

**(B)** Cytotoxicity assay of 5S- or TMG-treated HCCC-9810 cells. The cells were treated with 5S or TMG at various concentrations for 48 h.

Data were shown as the mean ± SD; statistical significance was determined by Student's t-tests (two-tailed, ***P < 0.001, ****P < 0.0001).

**Figure S5, related to Figure 2**

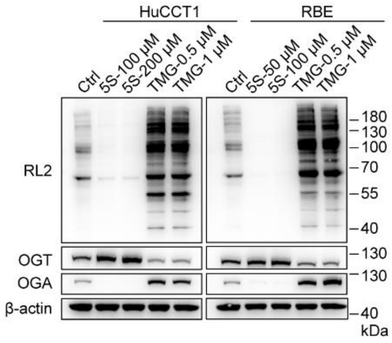


**Figure S5. *O*-GlcNAcylation can be adjusted in CCA cells with 5S or TMG, related to Figure 2.**

The analysis of the *O*-GlcNAcylated protein, OGT and OGA levels in 5S- or TMG-treated HuCCT1 and RBE cells by Western blot. Equal loading was confirmed using β-actin.

#### Figure S6, related to Figure 3

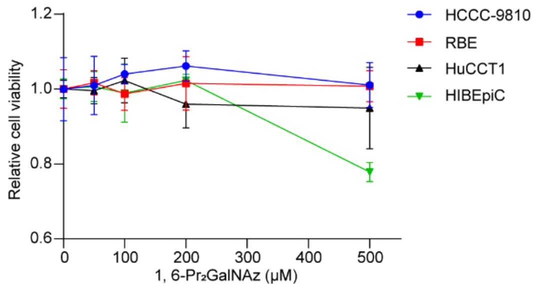


**Figure S6. Cytotoxicity of 1,6-Pr$_2$GalNAz in CCA cell lines and the Human Intrahepatic Biliary Epithelial Cell (HIBEpiC) control cell line by CCK-8 assays, related to Figure 3.**

1,6-Pr$_2$GalNAz, 1,6-di-*O*-propionyl-*N*-azidoacetylgalactosamine. Data were shown as the mean ± SD.

Figure S7, related to Figure 3

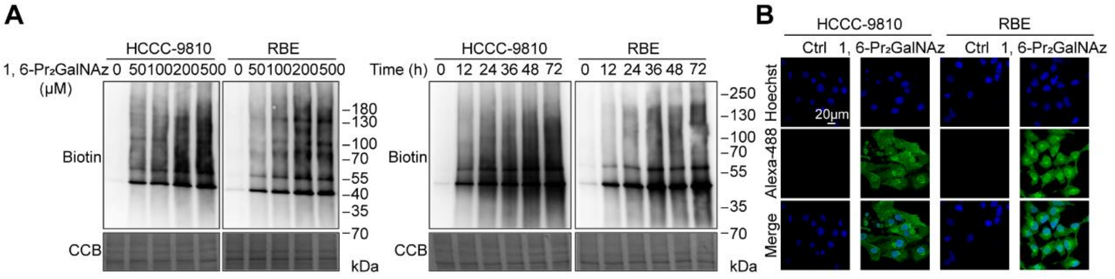


**Figure S7. The metabolic efficacy of 1,6-$Pr_2$GalNAz in HCCC-9810 and RBE cells, related to Figure 3.**

**(A)** Western blot analysis of HCCC-9810 and RBE cells treated with 1,6-$Pr_2$GalNAz at various concentrations for different times. The cell lysates were reacted with alkyne-biotin via copper(I)-catalyzed azide-alkyne cycloaddition (CuAAC), and blotted using anti-biotin. Equal loading was confirmed using Coomassie brilliant blue staining (CBB). **(B)** Confocal fluorescence imaging of HCCC-9810 and RBE cells treated with 1,6-$Pr_2$GalNAz at 0 or 200 μM for 48 h. The cells were washed, labeled with alkyne-AZDye-488, and analyzed. Scale bar: 20 μm.

#### Figure S8, related to Figure 3

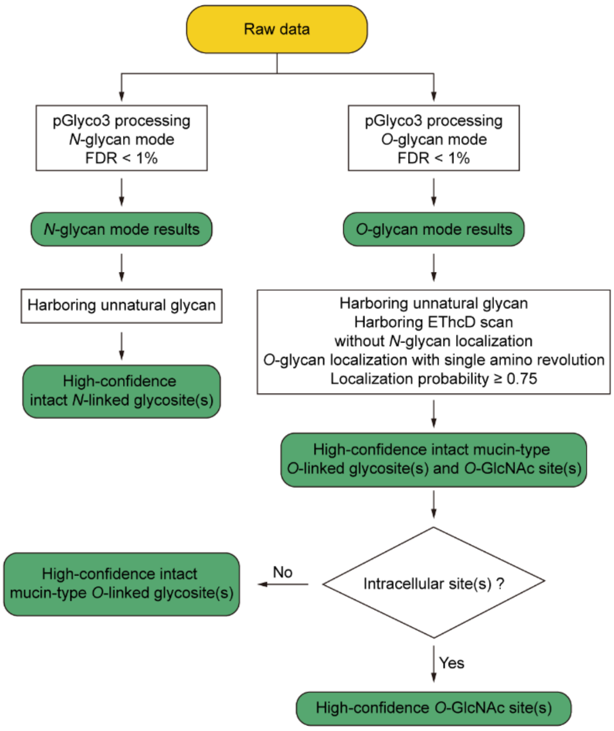


**Figure S8. Streamlined pipeline for screening high-confidence intact glycosites, related to Figure 3.**

#### Figure S9, related to Figure 3

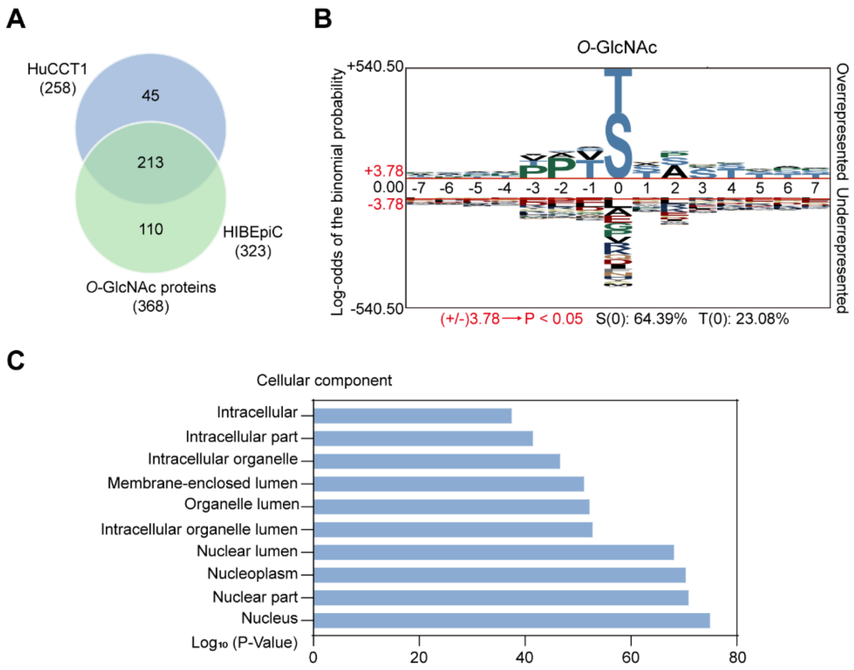


**Figure S9. Profiling of protein *O*-GlcNAcylation in HuCCT1 and HIBEpiC cells, related to Figure 3.**

**(A)** Overlap of the identified *O*-GlcNAcylated proteins between HuCCT1 and HIBEpiC cells.

**(B)** Motif analysis of *O*-GlcNAc sites identified in HuCCT1 and HIBEpiC cells using the pLogo. The log-odds binomial probability and adjusted P-value (adjusted by a conservative Bonferroni correction) were shown.

**(C)** Cellular component gene ontology (GO) terms of the identified *O*-GlcNAcylated proteins in HuCCT1 and HIBEpiC cells using the Database for Annotation, Visualization and Integrated Discovery (DAVID). The annotation terms and enrichment P-values (adjusted by a modified Fisher's exact test) were shown.

### Figure S10, related to Figure 3

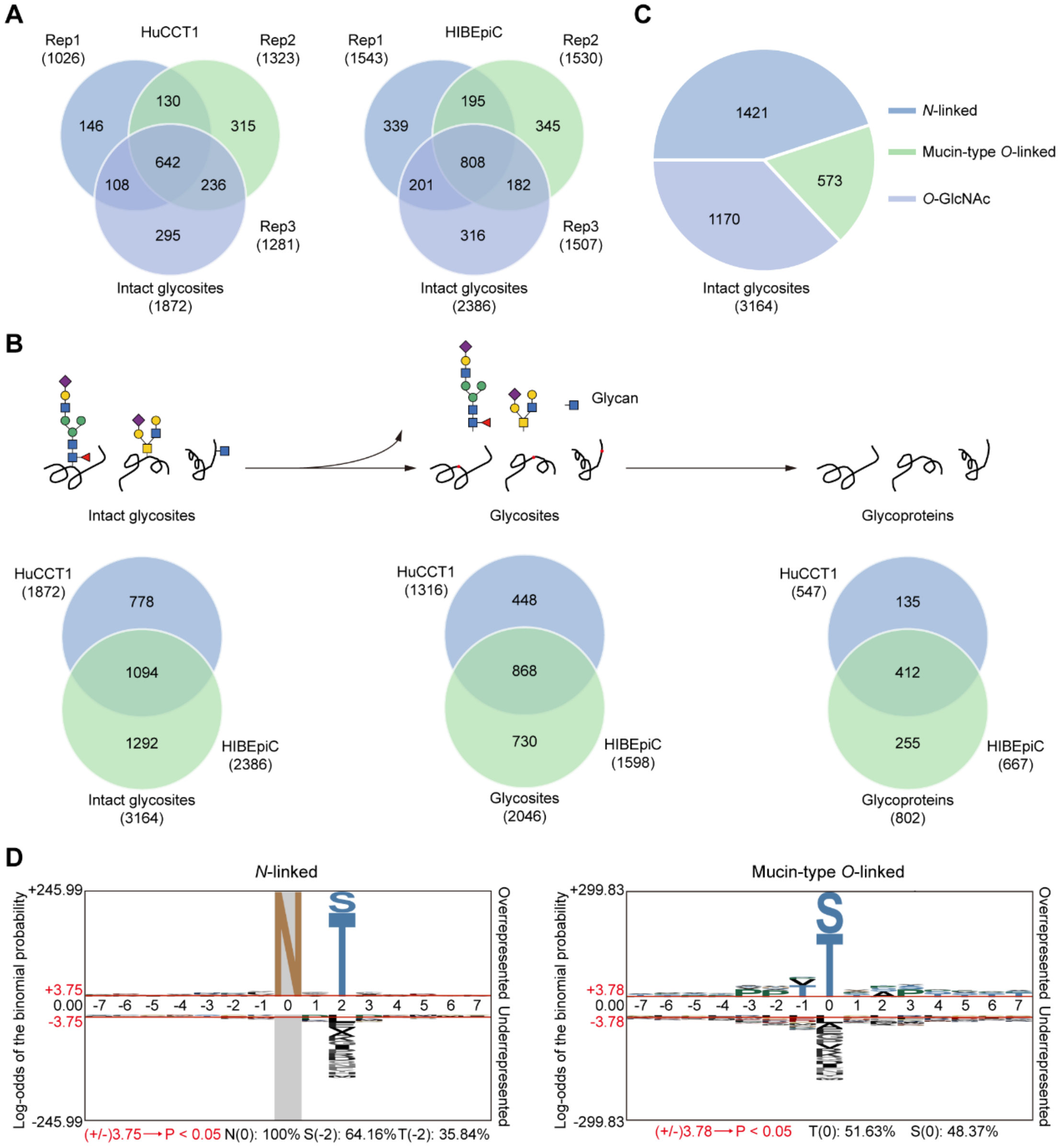

**Figure S10. Profiling of the intact glycosites in HuCCT1 and HIBEpiC cells, related to Figure 3.**

**(A)** Overlap of the identified intact glycosites in HuCCT1 or HIBEpiC cells between three replicates.

**(B)** Overlap of the identified intact glycosites, glycosites and glycoproteins between HuCCT1 and HIBEpiC cells.

**(C)** Total numbers and percentages of the intact *N*-linked glycosites, intact mucin-type *O*-linked glycosites, and intact *O*-GlcNAc sites identified in HuCCT1 and HIBEpiC cells.

**(D)** Motif analysis of the *N*-linked glycosites and mucin-type *O*-linked glycosites identified in HuCCT1 and HIBEpiC cells using the pLogo. The log-odds binomial probability and adjusted P-value (adjusted by a conservative Bonferroni correction) were shown.

**Figure S11, related to Figure 3**

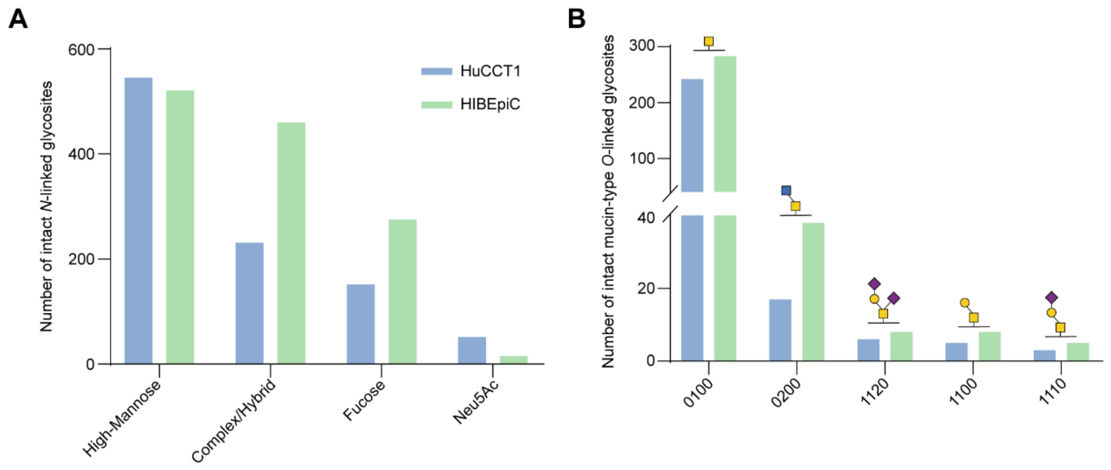


**Figure S11. Numbers of the intact glycosites with specific glycan in HuCCT1 and HIBEpiC cells, related to Figure 3.**

**(A)** Numbers of the intact *N*-linked glycosites with specific types (High-Mannose, Complex/Hybrid, Fucose and Neu5Ac) of *N*-glycans.

**(B)** Numbers of the intact mucin-type *O*-linked glycosites with specific *O*-glycans.

Neu5Ac, *N*-acetylneuraminic acid.

#### Figure S12, related to Figure 3

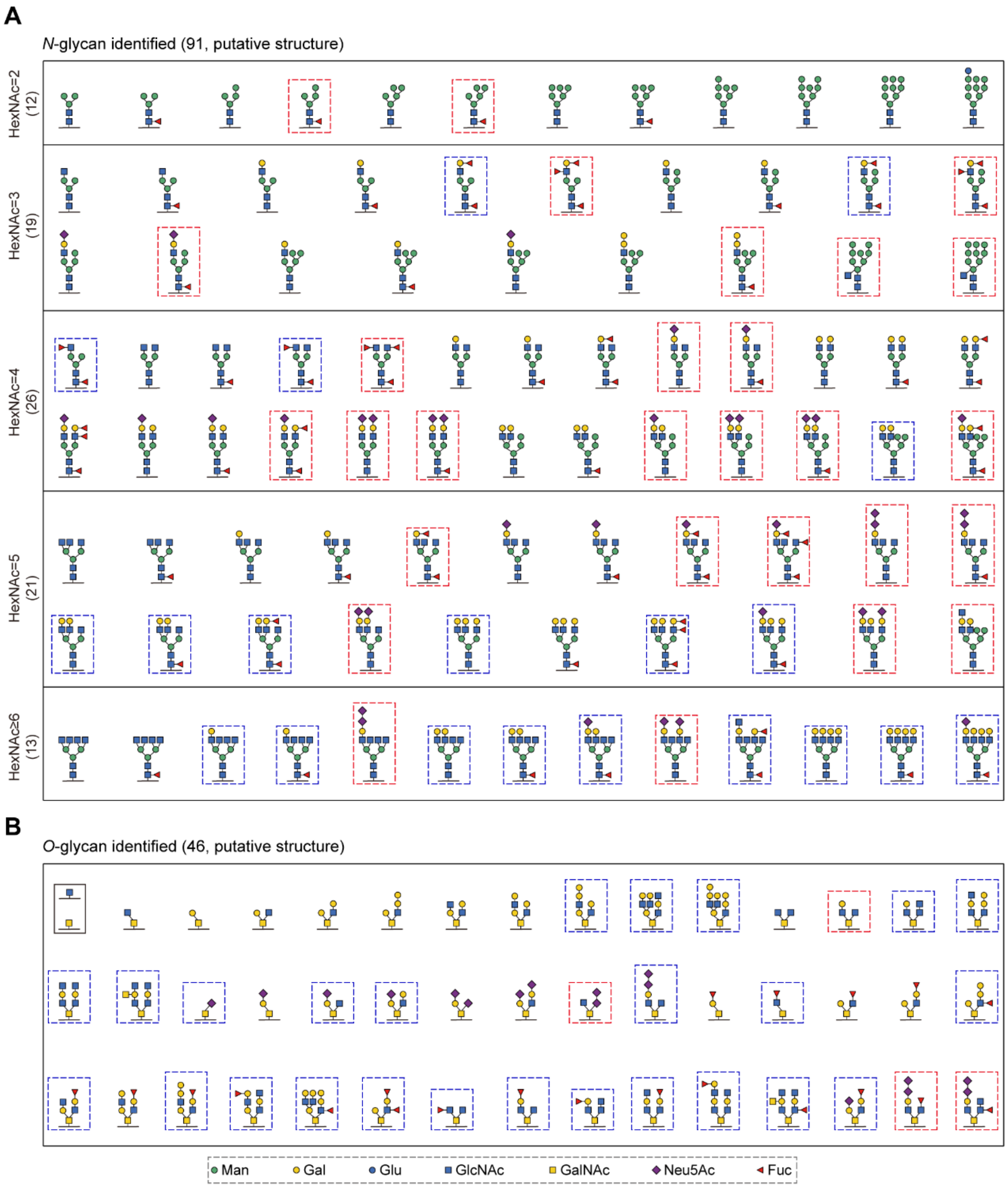

**Figure S12. Glycan compositions from the intact glycosites in HuCCT1 and HIBEpiC cells, related to Figure 3.**

**(A)** The putative glycan structure of 91 *N*-glycans was shown.

**(B)** The putative glycan structure of *O*-GlcNAc and 45 mucin-type *O*-glycans was shown.

The red or blue boxes were identified only in HuCCT1 or HIBEpiC cells, respectively. Man, mannose; Gal, galactose; Glu, glucose; GlcNAc, *N*-acetylglucosamine; GalNAc, *N*-acetylgalactosamine; Fuc, fucose.

#### Figure S13, related to Figure 3

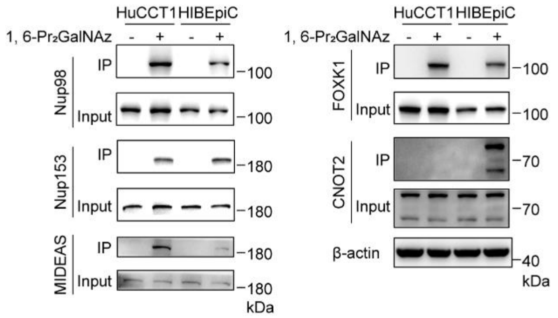


**Figure S13. Biochemical validation of identified *O*-GlcNAcylated proteins in HuCCT1 and HIBEpiC cells, related to Figure 3.**

Western blot analysis showing the *O*-GlcNAcylated candidates (Nup98, Nup153, MIDEAS, FOXK1 and CNOT2) in HuCCT1 and HIBEpiC cells. The cells incubated with 1,6-$Pr_2$GalNAz, lysed, reacted with alkyne-biotin, and captured by streptavidin beads. Equal loadings were confirmed using β-actin. IP, immunoprecipitation.

#### Figure S14, related to Figure 4

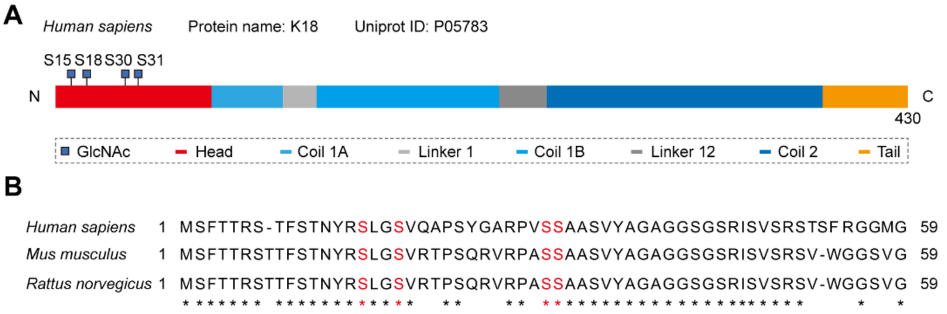


**Figure S14. *O*-GlcNAc sites of Keratin 18 (K18) identified in HuCCT1 and HIBEpiC cells, related to Figure 4.**

**(A)** Schematic showing the *O*-GlcNAc sites of K18.

**(B)** The *O*-GlcNAc sites of K18 are conserved across different species. *Human sapiens* (NP_001191439.1), *Mus musculus* (NP_001152847.1), *Rattus norvegicus* (NP_001093628.1).

Figure S15, related to Figure 5

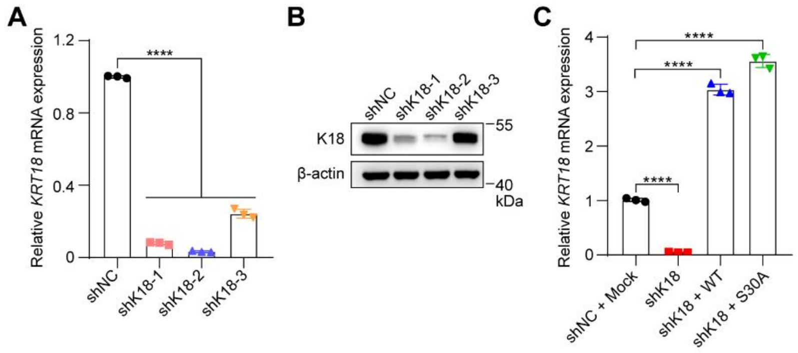


**Figure S15. Generation of HuCCT1 stable cell lines, related to Figure 5.**

**(A)** The *KRT18* mRNA expression of HuCCT1 stable cell lines with small hairpin RNA K18 knockdown (shK18-1~3) by qRT-PCR. Random small hairpin RNA (shNC) was used as a negative control.

**(B)** Western blot analysis of K18 in HuCCT1 stable cell lines with shK18-1~3. Equal loadings were confirmed using β-actin.

**(C)** qRT-PCR analysis of the *KRT18* mRNA expression in HuCCT1 stable cell lines with small hairpin RNA K18 knockdown (shK18) and re-expression of shK18-resistant FLAG-K18 wild-type (shK18 + WT) or FLAG-K18 S30A (shK18 + S30A). Random small hairpin RNA with an empty vector (shNC + Mock) was used as a negative control. Data were shown as the mean ± SD; statistical significance was determined by Student's t-tests (two-tailed, ****P < 0.0001).

#### Figure S16, related to Figure 5

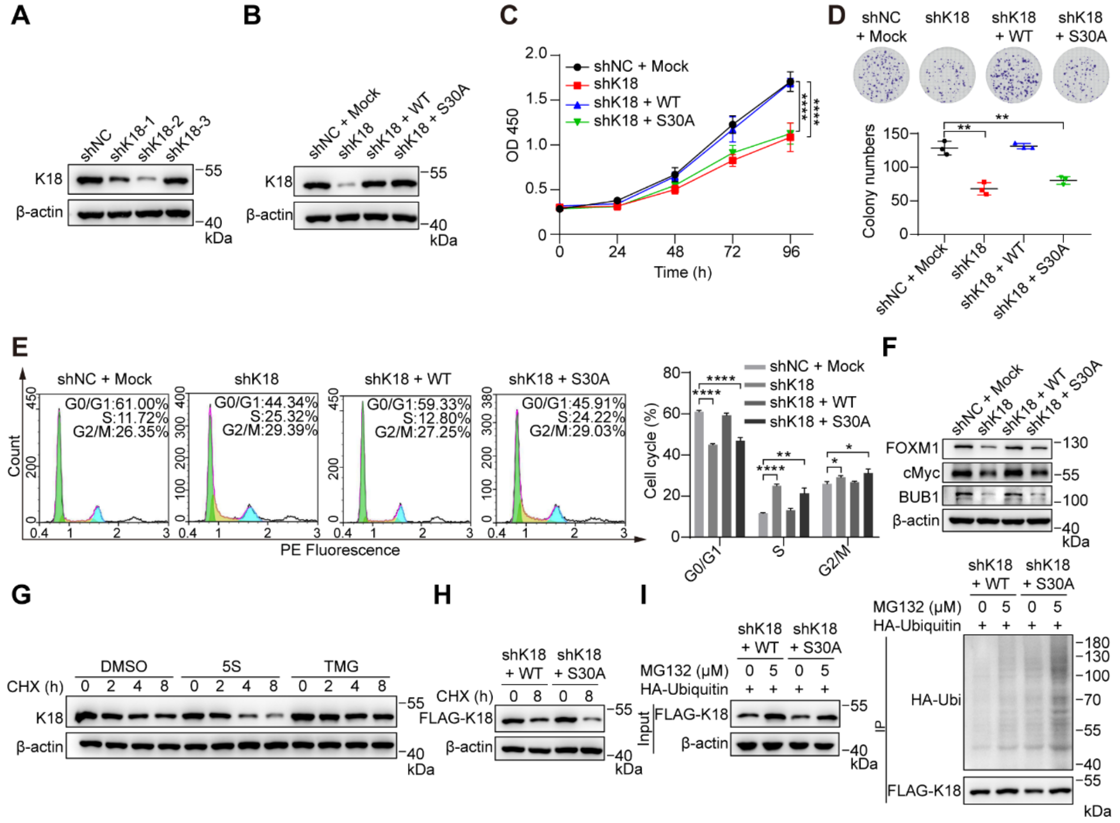

**Figure S16. *O*-GlcNAcylation of K18 promotes cell growth in RBE cells, related to Figure 5.**

**(A)** Western blot analysis of K18 in RBE stable cell lines with shK18-1~3.

**(B)** Western blot analysis of K18 in RBE stable cell lines with small hairpin RNA K18 knockdown (shK18) and re-expression of shK18-resistant FLAG-K18 wild-type (shK18 + WT) or FLAG-K18 S30A (shK18 + S30A). Random small hairpin RNA with an empty vector (shNC + Mock) was used as a negative control.

**(C)** CCK-8 analysis of RBE stable cell lines. Optical Density (OD) 450 was measured for cell viability.

**(D)** Clonogenic assay of cell proliferation in RBE stable cell lines. Colony numbers were quantitatively analyzed at the bottom.

**(E)** Cell cycle distribution assays of RBE stable cell lines. Histogram plot in flow cytometry indicated the percentage of cell populations in the G0/G1, S, or G2/M phase. Quantitative analysis was shown in the right panel.

**(F)** Cell cycle marker analysis of RBE stable cell lines by Western blot. Protein levels of FOXM1, cMyc (G1/S transition markers); BUB1(G2/M transition marker) were analyzed.

**(G)** Degradation analysis of K18 in RBE cells analyzed by Western blot. The cells were incubated with DMSO (vehicle), 200 μM 5S or 1 μM TMG for 48 h, followed by treatment with 10 μg/ml cycloheximide (CHX) for up to 8 h.

**(H)** Degradation analysis of K18 in RBE shK18 + WT or shK18 + S30A stable cell lines by Western blot. The cells were incubated with DMSO (vehicle) or 10 μg/ml CHX for 8 h.

**(I)** Ubiquitination analysis of K18 in RBE shK18 + WT or shK18 + S30A stable cell lines. The cells were transfected with HA-ubiquitin, incubated with 5 μM MG-132 (proteasome inhibitor) for 20 h, lysed and captured with anti-FLAG beads (left). Anti-HA blot demonstrated the ubiquitination of immunoprecipitated FLAG-K18 (right).

Equal loadings were confirmed using β-actin in all Western blot analyses. CCK-8, cell counting kit-8. Data were shown as the mean ± SD; statistical significance was determined by Student's t-tests (two-tailed, *P < 0.05, **P < 0.01, ****P < 0.0001).

**Figure S17, related to Figure 5**

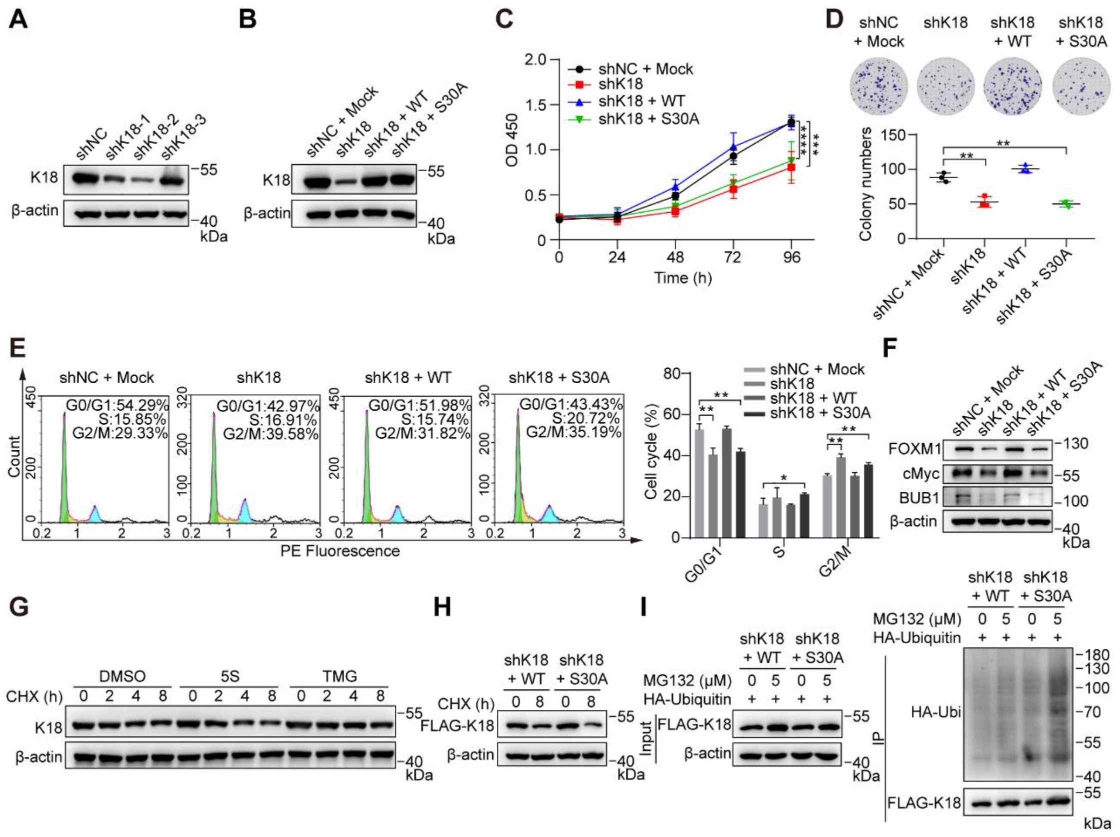

**Figure S17. *O*-GlcNAcylation of K18 promotes cell growth in HCCC-9810 cells, related to Figure 5.**

**(A)** Western blot analysis of K18 in HCCC-9810 stable cell lines with shK18-1~3.

**(B)** Western blot analysis of K18 in HCCC-9810 stable cell lines with small hairpin RNA K18 knockdown (shK18) and re-expression of shK18-resistant FLAG-K18 wild-type (shK18 + WT) or FLAG-K18 S30A (shK18 + S30A). Random small hairpin RNA with an empty vector (shNC + Mock) was used as a negative control.

**(C)** CCK-8 analysis of HCCC-9810 stable cell lines. Optical Density (OD) 450 was measured for cell viability.

**(D)** Clonogenic assay of cell proliferation in HCCC-9810 stable cell lines. Colony numbers were quantitatively analyzed at the bottom.

**(E)** Cell cycle distribution assays of HCCC-9810 stable cell lines. Histogram plot in flow cytometry indicated the percentage of cell populations in the G0/G1, S, or G2/M phase. Quantitative analysis was shown in the right panel.

**(F)** Cell cycle marker analysis of HCCC-9810 stable cell lines by Western blot. Protein levels of FOXM1, cMyc (G1/S transition markers); BUB1(G2/M transition marker) were analyzed.

**(G)** Degradation analysis of K18 in HCCC-9810 cells analyzed by Western blot. The cells were incubated with DMSO (vehicle), 200 μM 5S or 1 μM TMG for 48 h, followed by treatment with 10 μg/ml cycloheximide (CHX) for up to 8 h.

**(H)** Degradation analysis of K18 in HCCC-9810 shK18 + WT or shK18 + S30A stable cell lines by Western blot. The cells were incubated with DMSO (vehicle) or 10 μg/ml CHX for 8 h.

**(I)** Ubiquitination analysis of K18 in HCCC-9810 shK18 + WT or shK18 + S30A stable cell lines. The cells were transfected with HA-ubiquitin, incubated with 5 μM MG-132 (proteasome inhibitor) for 20 h, lysed and captured with anti-FLAG beads (left). Anti-HA blot demonstrated the ubiquitination of immunoprecipitated FLAG-K18 (right).

Equal loadings were confirmed using β-actin in all Western blot analyses. Data were shown as the mean ± SD; statistical significance was determined by Student's t-tests (two-tailed, $^{*}P < 0.05$, $^{**}P < 0.01$, $^{***}P < 0.001$, $^{****}P < 0.0001$).

**Figure S18, related to Figure 5**

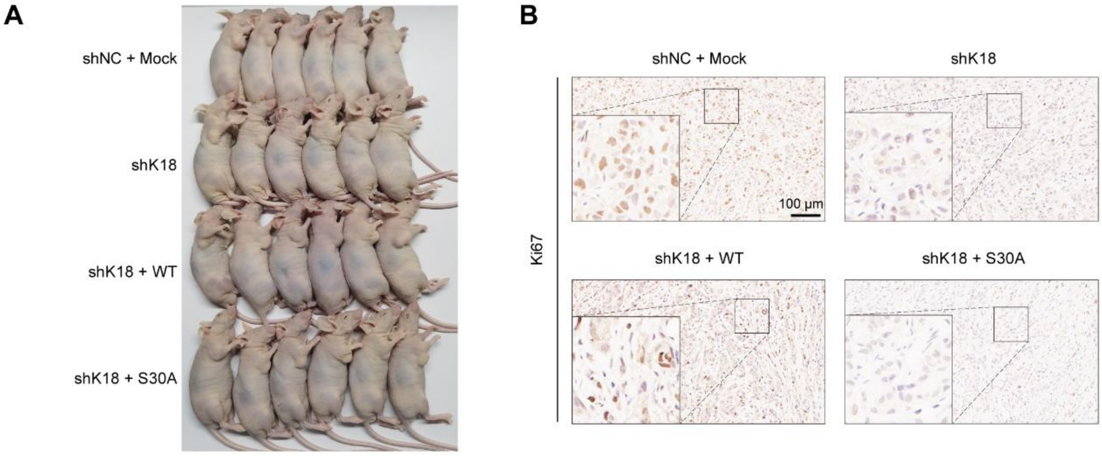


**Figure S18. Tumor formation in nude mice, related to Figure 5.**

**(A)** Photograph of tumors generated by xenograft HuCCT1 stable cell lines.

**(B)** Representative images of Ki67 staining in xenograft tumors through immunohistochemistry (IHC). Scale bar: 100 μm.

**Figure S19, related to Figure 5O**

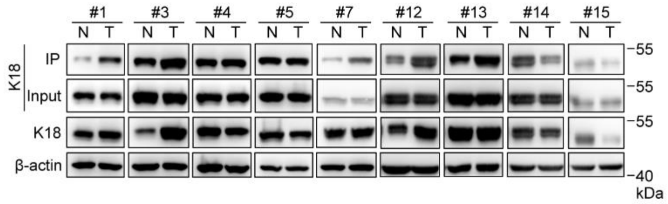


**Figure S19. K18 and its *O*-GlcNAcylation are dysregulated in CCA, related to Figure 5O.**

Representative images of K18 and its O-GlcNAcylation levels from nine pairs of CCA tumor tissues (T) and adjacent normal tissues (N) by Western blot analysis. Equal loadings were confirmed using β-actin.

**Figure S20, related to Figure 6**

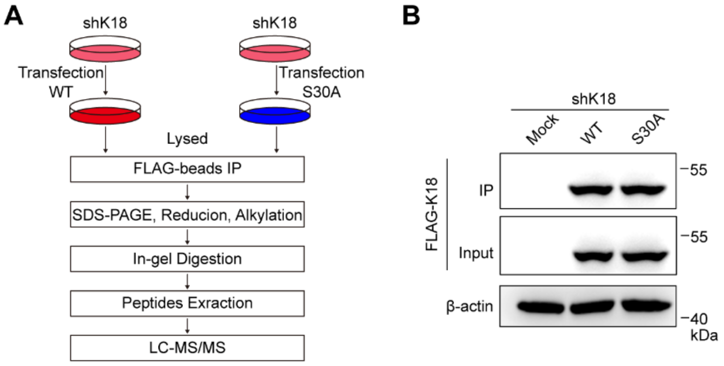


**Figure S20. The analysis of interacting proteins of K18 protein, related to Figure 6.**

**(A)** Workflow of the analysis of interacting proteins with K18 in HuCCT1 shK18 cells transfected with FLAG-K18$^{WT}$ or FLAG-K18$^{S30A}$ by liquid chromatography-tandem mass spectrometry (LC-MS/MS).

**(B)** Western blot and immunoprecipitation analysis showing the FLAG-K18 protein levels in HuCCT1 shK18 cells transfected with FLAG-K18$^{WT}$ or FLAG-K18$^{S30A}$. Equal loadings were confirmed using β-actin.

**Figure S21, related to Figure 6**

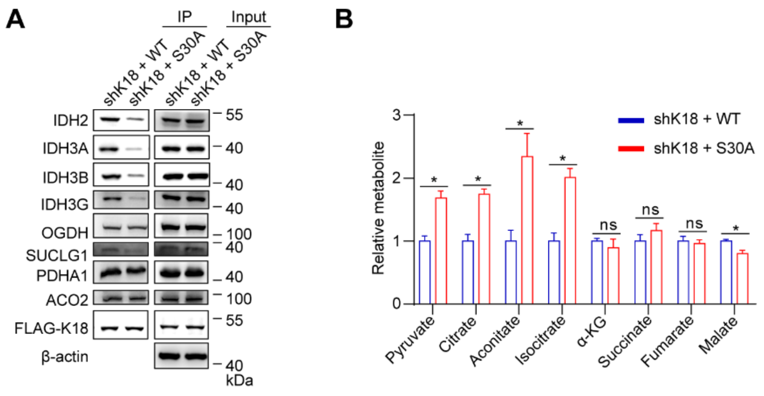


**Figure S21. *O*-GlcNAcylation of K18 regulates the metabolism of TCA cycle, related to Figure 6.**

**(A)** The analysis of enzymes in the TCA cycle interacting with K18 of RBE stable cell lines (shK18 + WT and shK18 + S30A). Protein levels of IDH2, IDH3A, IDH3B, IDH3G, OGDH, SUCLG1, PDHA1, ACO2 and FLAG-K18 were analyzed by Western blot and immunoprecipitation analysis. Equal loadings were confirmed using β-actin.

**(B)** Relative abundance of metabolites of the TCA cycle in RBE stable cell lines (shK18 + WT and shK18 + S30A). Data were shown as the mean ± SD; statistical significance was determined by Student's t-tests (two-tailed, $^*P < 0.05$, ns, not significant).
